# PER2- and state-dependent transcriptional programs gate neural stem cell proliferation with niche-specific circadian autonomy

**DOI:** 10.64898/2026.09.14.751519

**Authors:** Pedro O. Brum, Ava Abdi, Federico Scaramuzza, Tatjana Kepcija, Michael H. Hastings, Andrew S.I. Loudon, Kristin Tessmar-Raible, Noelia Urbán

## Abstract

Adult neural stem cells (NSCs) in the mouse brain are predominantly quiescent, with activation tightly regulated to balance neurogenesis and stem cell maintenance. Circadian clocks temporally organize core cellular processes, potentially gating NSC activation. We examined adult NSC temporal dynamics across the day and observed rhythmic expression of core circadian clock components BMAL1 and PER2 in both mammalian niches, the subgranular zone (SGZ) of the dentate gyrus and the subventricular zone (SVZ). While cell cultures derived from both niches show BMAL1 and PER2 protein, only SGZ-derived NSCs exhibit synchronized self-sustained core clock oscillations across the population. Comparative analyses of WT and *Per2* knockout NSCs revealed state-specific, circadian clock-dependent oscillatory transcriptional programs. This identified ASCL1 and CCND1 as candidate regulators of cell-cycle coordination, with daytime accumulation preceding a nighttime S-phase peak. The temporal control of S-phase entry in NSCs was abolished in *Per2* knockout mice. Together, our findings reveal state-specific, PER2-dependent circadian regulation of adult NSCs in both neurogenic niches, and a specific dependence on external stimuli for synchronization in the SVZ.

**Highlights:**

- Core circadian clock proteins BMAL1 and PER2 display diel rhythmicity in quiescent and active adult NSCs of SGZ and SVZ.
- SGZ-derived but not SVZ-derived NSCs maintain synchronicity of their circadian clock outside of the niche.
- PER2-dependent oscillations govern distinct transcriptional programs in active and quiescent NSCs of the SGZ.
- Daytime accumulation of circadian targets ASCL1 and CCND1 precedes the nocturnal accumulation of S-phase NSCs in the SGZ.
- *Per2* is essential for the temporal gating of S-phase entry in both niches.

## 1. Introduction

Organisms respond and adapt to prominent daily oscillations in their environment, such as light and temperature, and consequently, many biological processes exhibit rhythmicity. This rhythmicity occurs either as a direct response to the changing environment or as an output of endogenous oscillators. Among the latter, the endogenous circadian clock coordinates cellular programs that occur in a ∼24-h period. This temporal coordination is most commonly mediated directly and indirectly by interlocked, self-sustained transcription-translation feedback loops (TTFL) that constitute the core molecular clockwork^1^.

The transcription factors CLOCK and BMAL1 form a heterodimer that binds E-box elements to activate rhythmic gene expression, forming the primary positive loop of the mammalian core circadian clock. This activation is counterbalanced by the repressors PER (*Per1*, *Per2*, *Per3*) and CRY (*Cry1* and *Cry2)*: following their expression, PER/CRY complexes are phosphorylated, triggering their translocation to the nucleus, where they inhibit CLOCK/BMAL1-dependent transcription, thereby closing the primary negative feedback loop^1^. Indeed, comprehensive transcriptomic^2^ and proteomic^3^ atlases have revealed that a vast array of biological processes are temporally segregated by the circadian clock to optimize cell and tissue functions in mammals, such as cell-cycle progression and proliferation^4,5^ and cell-fate decisions^6,7^.

Adult stem cells provide a stable reservoir for the continuous renewal of specialized cells across diverse tissues. Rhythmic, circadian clock-controlled proliferation is a common feature of adult stem cell niches, with genetic ablation of core clock components (*Per1*, *Per2*, *Bmal1*) altering the timing of cell-cycle entry, cell-fate decisions, and/or differentiation across multiple peripheral tissues ^8,9^. Circadian gating of proliferation is thought to coordinate cell-cycle entry with periods of low metabolic demand, minimizing replication-associated DNA damage and preserving long-term stem cell pool integrity^9^. In the mammalian central nervous system stem cells reside in two specialized niches: the subventricular zone (SVZ) of the lateral ventricles and the subgranular zone (SGZ) of the dentate gyrus^10^. Most adult neural stem cells (NSCs) remain in a quiescent state, reversibly exiting the cell cycle to preserve their long-term neurogenic potential ^11^. Upon activation, NSCs re-enter the cell cycle to self-renew and/or differentiate and integrate into pre-existing circuits^12^. This is tightly regulated by intrinsic and extrinsic cues. The limited self-renewal capacity of NSCs couples activation to differentiation, resulting in adult stem cell depletion over time^13^.

Diel rhythms encompass a full day-night cycle, irrespective of endogenous clock or environmental control. Increasing evidence suggests that diel activation may be essential for preserving NSC proliferation dynamics and cell-fate^14–17^. However, results regarding NSC circadian clock rhythmicity have remained ambiguous. This ambiguity emerges from several overlapping methodological and biological challenges: the lack of cross-niche comparisons, insufficient cell-type resolution and unresolved clock dynamics between active and quiescent NSC states, as well as the unclear contributions of signals such as melatonin.

Higher SVZ NSC division rates during the daytime have been suggested to be mediated directly by the day/night cycle via melatonin signalling^16^. However, most commonly used laboratory mouse strains are melatonin-deficient^18,19^, yet diel regulation of adult NSCs is robustly observed in the SGZ^15,17^. Moreover, rhythmic BrdU incorporation in the SGZ persists under constant darkness^15^, arguing that diel control of adult neurogenesis cannot be solely driven by melatonin or direct light conditions. Instead, PER2 or BMAL1 knockouts lead to increased overall proliferation and/or loss of diel organization of cell divisions^15^. This implicates the circadian clock machinery as a melatonin-independent regulator of SGZ NSCs. Whether a similar circadian clock-driven logic governs diel NSC activation in the SVZ, and how the two niches compare, remains to be elucidated.

Another challenge concerns cell-type resolution. Studies reporting diel changes in PER2 across the SGZ (mPer2-DsRed^15^) and SVZ (PER2::LUC^20^) relied on bulk cell counts or whole-tissue bioluminescence, lacking cell-type specificity. Previous reports attempted to address this using Nestin-GFP mice, for NSCs specificity^17^. However, the percentage of BMAL1 positive cells displayed only a shallow trough which overlaps with the previously reported trough of mPer2-DsRed cell numbers ^15,17^. This overlap is difficult to reconcile with the canonical function of the circadian clock, highlighting the need for quantifying intensity in individual cells rather than counting percentages of positive cells. Furthermore, while comparisons between proliferating and non-proliferating cells have suggested lower levels of core clock components in active compared to quiescent SGZ NSCs^15^, the distinction between cell states has not been accounted for with temporal resolution.

Aiming to bridge these gaps, we demonstrate that NSCs in both the SVZ and SGZ possess a functional molecular clock that temporally organizes cell-cycle progression *in vivo*. However, only SGZ-derived NSCs maintain synchronized circadian clock function *in vitro*, suggesting distinct strengths of the cellular circadian clocks between niches. Using circadian transcriptomics, we identified PER2-dependent, state-specific oscillations in SGZ NSCs. In active NSCs these are driven by energy metabolism, cell cycle and neurogenic genes, whereas in quiescent NSCs these are comprised of biosynthetic and lysosomal genes. Based on the transcriptomic findings, we identified Ccnd1 and Ascl1 as PER2-dependent likely effectors of circadian-controlled NSCs activation. This is supported by their daytime protein accumulation preceding a nighttime peak in S-phase cells. The PER2-dependent circadian regulation of the cell cycle in both niches, together with their different intrinsic circadian clock strength points to differential dependence on external signals to coordinate rhythmicity in the two mammalian neurogenic niches.

## 2. Results

### 2.1 Rhythmic expression of core clock components in adult NSCs of the SVZ and SGZ

As outlined above, previous analyses have offered contrasting conclusions on the role of the circadian clock and its components in the two different mammalian adult NSC niches. To characterize the circadian clock within NSCs of both the SGZ and SVZ, we monitored a core component of the repressive loop, PER2, alongside the positive element, BMAL1. By quantifying their relative nuclear intensity levels in morphologically, phenotypically and cell-state identified NSCs, we established a quantitative readout over time of the intracellular circadian clock machinery.

We sampled PER2::VENUS reporter mice^21^ every 4 h from ZT0 to ZT20 (ZT: zeitgeber time, i.e. under light-dark entrainment). VENUS staining was used as a readout of PER2 protein levels. The impact of VENUS reporter fusion was thoroughly assessed and proven to reflect physiological PER2 kinetics previously^21^. BMAL1 levels were measured using an antibody against the endogenous protein^22^.

Adult NSCs were identified based on expression of GFAP and Sox2 together with location and morphology^23–25^. Adult NSCs display robust PER2::VENUS diel oscillations, peaking between ZT12 and ZT16 (Fig. 1E-F). By contrast, BMAL1 peaks in antiphase to PER2 at ZT8 (Fig. 1I–J). Quiescent and active NSCs were further distinguished utilizing markers associated to proliferation, MCM2 (Supp. Fig. 1-2) and Ki67^25^ (Supp. Fig. 3-4). Both active and quiescent NSCs in the SGZ and SVZ express PER2::VENUS and BMAL1. Quiescent NSCs display the same dynamics observed at the population level. In active NSCs, PER2::VENUS also showed the same dynamics (Supp. Fig. 1-2), however, BMAL1 levels remained constantly low in the SGZ (Supp. Fig. 3) and SVZ (Supp. Fig. 4). Despite these differences in BMAL1 levels, the overall relationship between BMAL1 and PER2 still indicates an intact, functional circadian oscillator within the NSCs of both niches and in both states.

**Figure 1:**
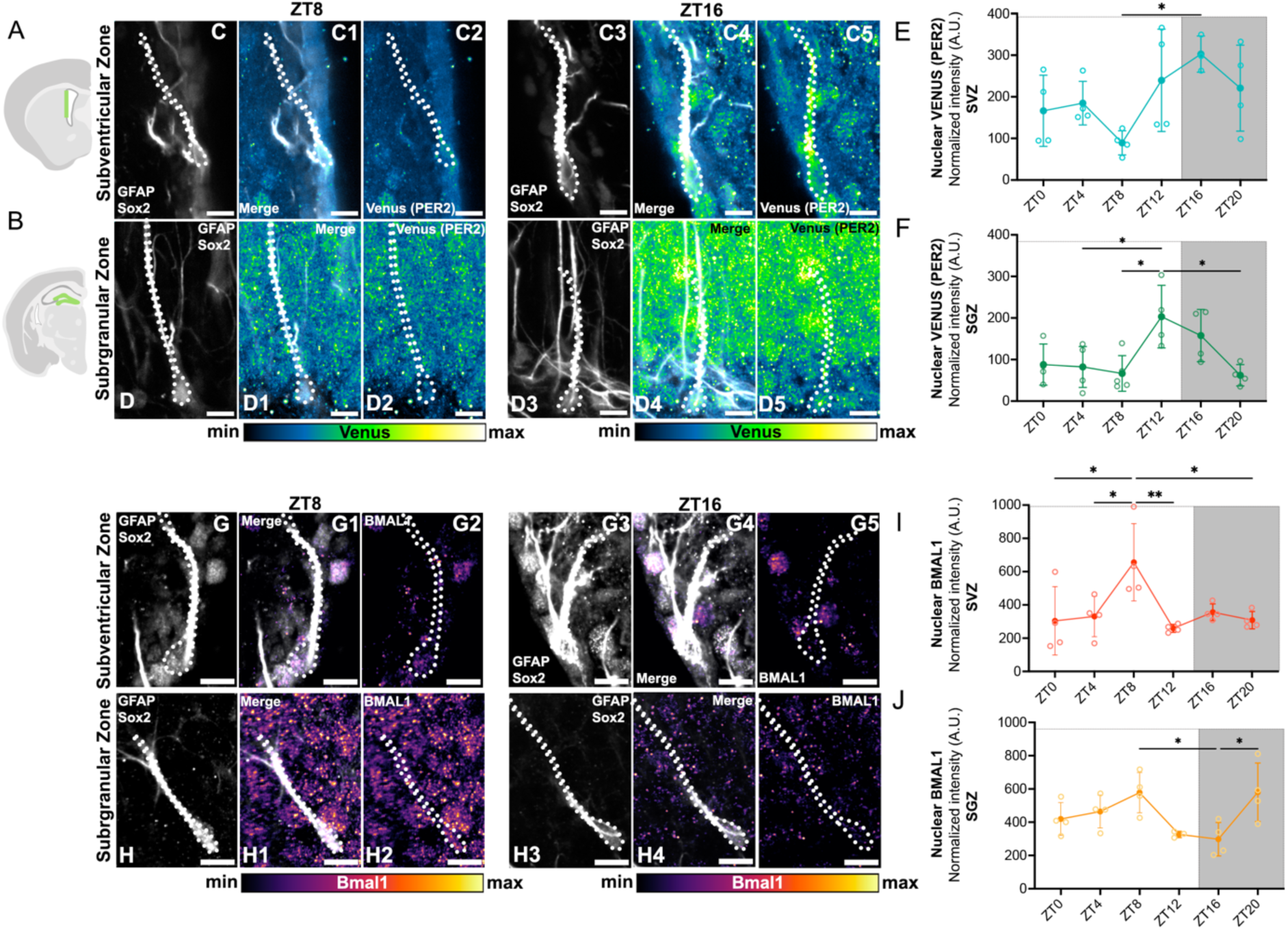
Diel expression of PER2 and BMAL1 in NSCs of the SGZ and SVZ suggests functional circadian clock activity. Brain sections containing the (**A**) SVZ and (**B**) SGZ were used for immunostaining; quantifications were focused within the highlighted green areas. (**C**-**C5**, **D**-**D5**, **G**-**G5**, **H**-**H5**) Nuclear fluorescence intensity for PER2::VENUS and BMAL1 in NSCs. Scale bar = 10µm. (**E**-**F**) Peak PER2::VENUS intensity was observed at ZT16 for both SVZ and SGZ NSCs; conversely, (**I**-**J**) peak BMAL1 levels were observed at ZT8 in both niches (One-way ANOVA followed by Tukey’s *post hoc* test; * *p* < 0.05, ** *p* < 0.01, *** *p* < 0.001). Open circles represent individual replicates; solid circles and error bars represent mean ± SD.

### 2.2 Niche-specific differences in circadian clock strength *in vitro*

To determine whether the circadian rhythms we observed *in vivo* are cell autonomously generated and/or synchronized (i.e. entrained), we derived adult NSC cultures from the SGZ and SVZ of PER2::VENUS mice. We sampled NSCs in active and quiescent conditions^26^ every 4 h over 36 h and measured PER2 and BMAL1 levels through immunostaining (Fig. 2). Active SGZ-derived cells exhibited robust oscillations for PER2::VENUS (Fig. 2B-C, cosinor, period = 24 h, *p* < 0.05), while quiescent SGZ-derived cells displayed rhythmicity for BMAL1 (Fig 2 D-E, cosinor, period = 24 h, *p* < 0.05). Active or quiescent SVZ-derived NSCs, on the other hand, failed to exhibit rhythmicity for either marker (cosinor, period = 24 h, *p* > 0.05; Fig. 2 F–I).

**Figure 2:**
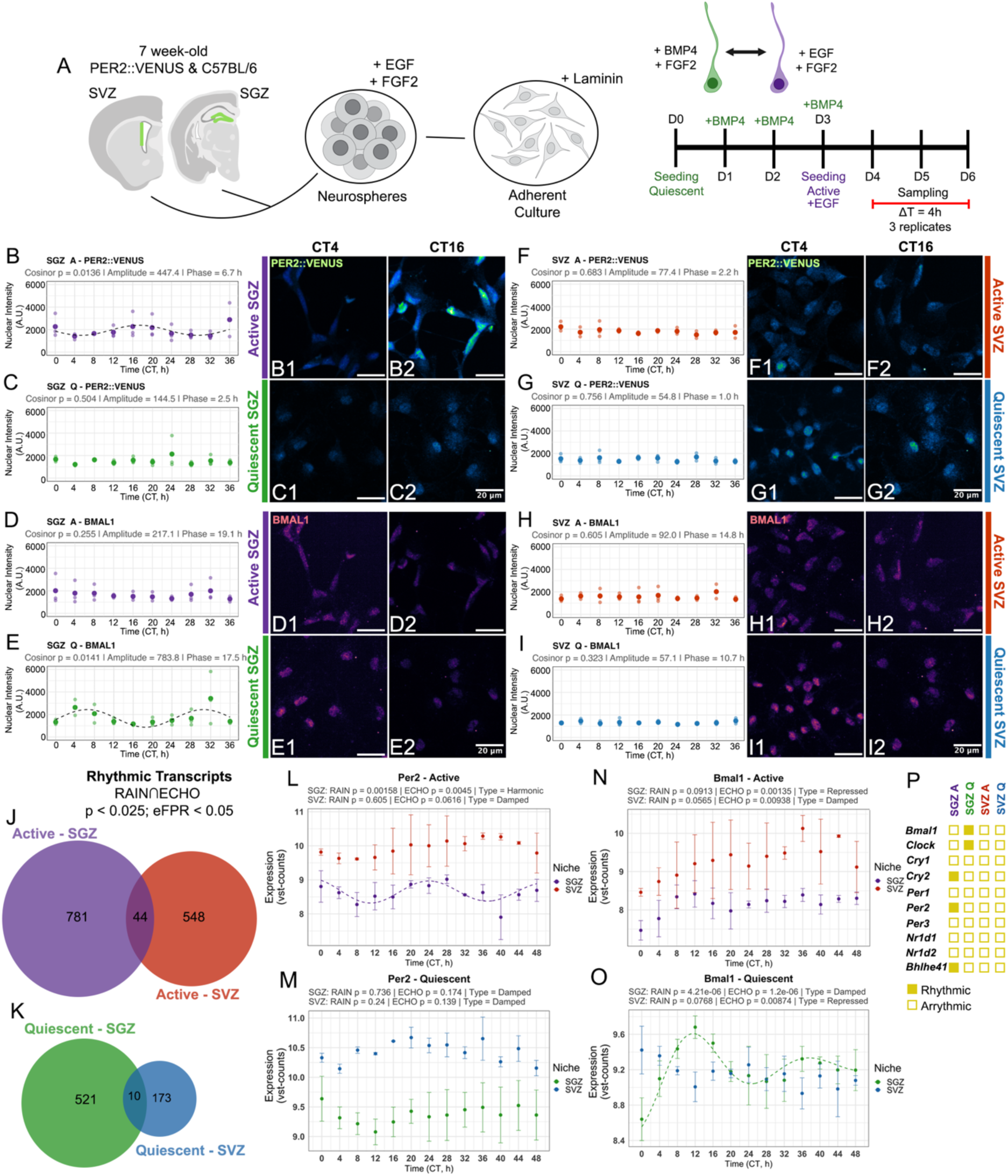
Niche-dependent autonomous circadian oscillations in adult neural stem cells. **(A)** Schematic representation of the experimental design. NSCs derived from the SVZ and SGZ of 7-week-old C57BL/6J or PER2::VENUS mice were maintained in either quiescent (BMP4 and FGF2) or active (EGF and FGF2) conditions. For protein analysis, cells were sampled every 4 h over 36 h, immunostained for GFP (PER2::VENUS) and BMAL1, and analyzed for nuclear fluorescence intensity. For transcriptomics, cells were sampled every 4 h over 48 h. **(B–I)** Temporal profiles of PER2::VENUS and BMAL1 nuclear fluorescence intensity in SGZ and SVZ NSCs. Traces are shown for the SGZ (**B–E**) and SVZ (**F–I**), alongside representative images of anti-phase timepoints. In all protein plots, lighter colored dots represent individual replicates, while darker dots indicate the mean for each time point, and cosinor fit curves (24 h period) are overlaid only for datasets exhibiting significant rhythmicity (*p* < 0.05). For transcriptomic analysis of circadian mRNA oscillations, rhythmic transcripts were identified using a consensus approach in RAIN and ECHO (*p* < 0.025 for both), filtering for Harmonic, Damped, or Forced fits and excluding Repressed or Overexpressed traces. **(J–K)** Venn diagrams display the number of shared and niche-specific rhythmic transcripts passing a permutation-based false positive rate (FPR < 5%) in active (**J**) and quiescent (**K**) states. **(L–O)** Temporal mRNA expression profiles of *Per2* and *Bmal1* in active and quiescent NSCs derived from the SGZ and SVZ. ECHO fits are displayed only for significantly rhythmic transcripts, and error bars represent SD (n=3). **(P)** Schematic representation of core clock network rhythmicity. Solid boxes denote transcripts identified as significantly rhythmic, whereas open boxes represent arrhythmic transcripts across the indicated niches and activation states (detailed temporal traces plots are provided in Supp. Fig. 5).

The circadian expression of clock components at the protein level in a bulk analysis suggests the presence of a functional, synchronized cell-autonomous circadian oscillator in SGZ NSCs. SVZ NSCs lost synchronized oscillations *in vitro*, highlighting profound niche-specific differences in circadian clock strength. Altogether, this suggests that SVZ NSCs rely more heavily on extrinsic cues than SGZ NSCs.

Next, to obtain a global view of circadian rhythmic transcripts in NSCs *in vitro*, we performed bulk RNA sequencing collected every 4 h over a 48-h period in active and quiescent conditions (Fig. 2 A). Rhythmicity was assessed using two independent R packages, ECHO^27^ and RAIN^28^, intersecting the results. Both methods complemented each other: RAIN detects monotonic slopes irrespective of waveform shape, while ECHO identifies harmonic oscillations with a damping factor. This is noteworthy, as *in vitro* transcriptomic datasets have been shown to exhibit a substantially higher proportion of damped oscillations compared to *in vivo* conditions, which may go undetected by fixed-amplitude methods^27^. Transcripts were considered rhythmic only when the *p*-value was < 0.025 in both algorithms. Additionally, an empirically determined false positive rate (eFPR) based on permutation tests of the datasets was applied as an additional filter with a cutoff of eFPR < 0.05 (see methods). By these highly stringent criteria we aimed to obtain only those transcripts that were most reliably rhythmic.

We first checked the rhythmicity of core clock machinery genes in both niches. SGZ NSCs exhibited a state-dependent core circadian clock expression: active cells predominantly displayed rhythmicity in negative-loop transcripts (*Per2, Cry1, Cry2, Bhlhe41*), whereas quiescent cells showed rhythmicity in positive-loop transcripts (*Bmal1, Clock*) (Fig. 2 L–P, Supp. Fig. 5 C-R). The cycling of *Clock* is particularly noteworthy, as its transcript is constitutively expressed in the suprachiasmatic nucleus. In SVZ-derived cultures, none of the circadian clock components showed a circadian expression pattern at the mRNA level (Fig. 2 P, Supp. Fig. 5).

Globally, active SGZ-NSCs exhibited 825 rhythmic transcripts. In active SVZ-NSCs, we identified 592 rhythmic transcripts, of which 44 were shared between active cells from the two regions (Fig. 2 J). While there were no enrichments among these shared rhythmic transcripts, 10 of these transcripts are related to cell cycle regulation and proliferation (*Tob2, Zfp521, Pard6g, Cdc42ep4, Mlx, Slc29a3, Stk38l, Tfip11, Nacc2,* and *Rrs1)* (Supp. Table 2). In the distinct rhythmic transcripts, active NSCs from both niches showed enrichment of functionally related pathways associated with RNA metabolism, ribosome biogenesis and regulation of cellular metabolic processes (Supp. Table 3).

Quiescent NSCs displayed an overall lower number of rhythmic transcripts than active NSCs: 531 in the SGZ and only 183 in the SVZ, with 10 shared between regions (Fig. 2K). Among these shared transcripts, 3 are involved in signaling regulation (*Ter2ip*, *Zc3h12c*, *Zcchc3*) (Supp. Table 2). In the distinct rhythmic transcripts, quiescent NSCs from both niches displayed enrichment in pathways associated with autophagy–lysosomal function, cellular organization/localization, RNA metabolism, and DNA repair (Supp. Table 3).

54 transcripts were shared between active and quiescent SGZ-NSCs and 8 transcripts were shared between conditions in SVZ-NSCs (Supp. Fig. 5A-B). Interestingly, the shared transcripts in SGZ-derived samples were highly enriched for sterol biosynthesis, which could be relevant in the context of NSC homeostasis (Supp. Table 3).

These state-and niche-specific protein and mRNA circadian patterns show that i) *in vitro* cultured NSCs from both niches are phase-synchronized, ii) core circadian clock synchronization is stronger in SGZ-NSCs than in SVZ-NSCs and iii) different transcriptional loops dominate in active versus quiescent NSCs.

### 2.3 PER2-dependent rhythmic transcription in SGZ-derived NSCs

Noting the autonomously synchronized circadian clock transcript and protein rhythmicity in SGZ-NSCs, we next asked which rhythmic transcriptional programs depend on the core circadian clock gene *Per2* in these cells. To address this, we sampled NSC cultures derived from *Per2* KO (P2KO) mice^29^ using the same experimental design described above (Fig. 2 A). Overall differential expression analysis displayed no alteration in the stemness profile upon deletion of *Per2* (Supp. Fig. 6).

The majority of transcripts (90,06% in active and 96,9% in quiescent conditions) lost rhythmicity in P2KO NSCs, showing a strong dependence of NSCs circadian rhythmicity on *Per2* and hence the core circadian clock (Fig. 3 B, C). PER2 deletion also led to a partial rewiring of the clock, with distinct components detected between WT and P2KO samples. In active P2KO NSCs, *Cry1* and *Bmal1* gained rhythmicity. In quiescent P2KO NSCs, the compensatory activator *Npas2*, lowly expressed or undetected in WT samples, gained rhythmicity alongside the repressor *Per3*, suggesting a state-specific compensatory response to PER2 loss (Fig. 3 D and Supp. Fig. 7). Such re-wiring of the circadian network might be interesting from the oscillator plasticity and evolutionary perspective, but have no correspondence under normal conditions. Hence, we decided to focus here on those transcripts that lost circadian rhythmicity in P2KO.

**Figure 3:**
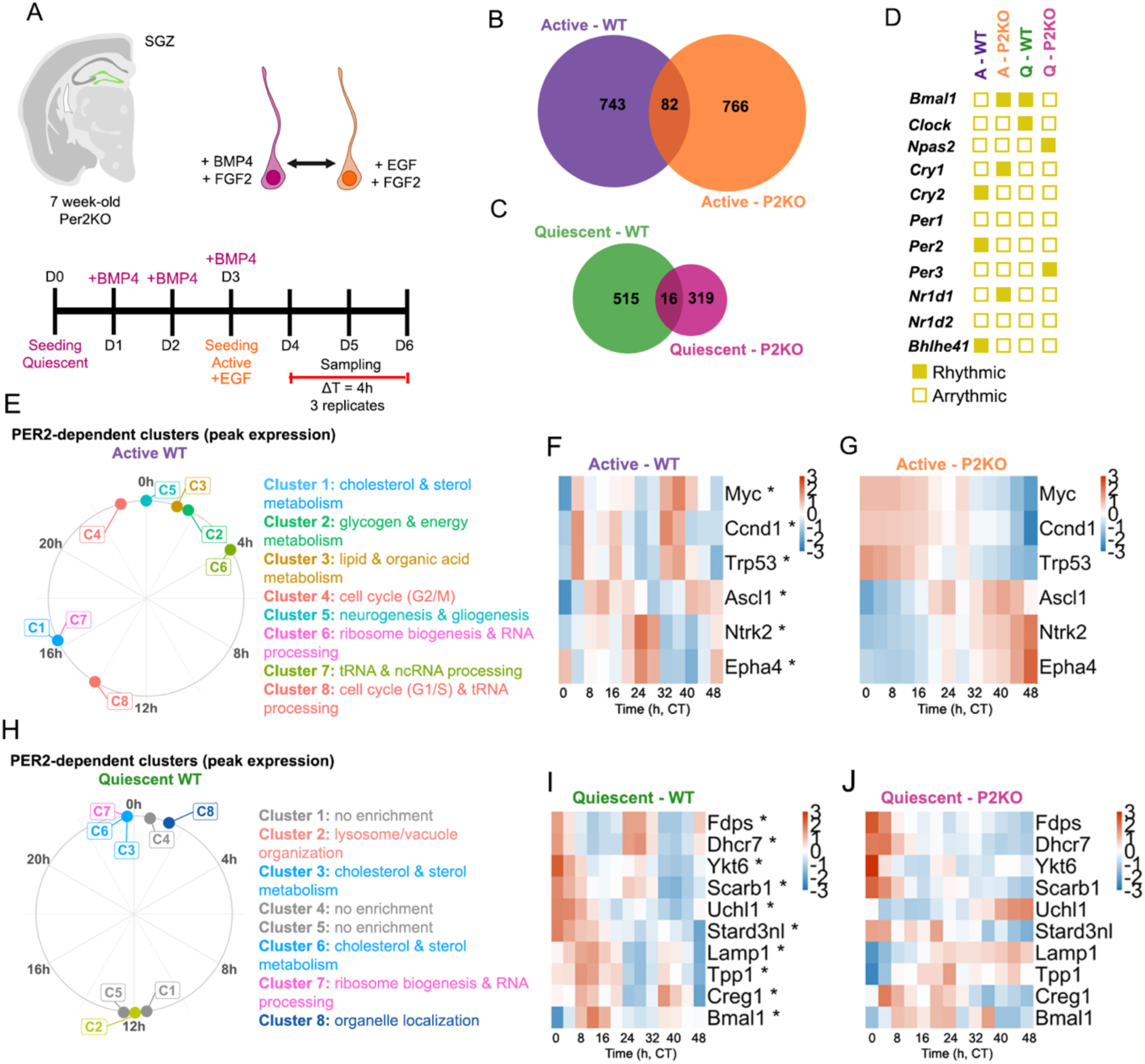
**(A)** SGZ NSCs were derived from 7 week-old *per2* KO mouse and circadian transcriptomics was performed as previously described (48h, ΔT = 4h, n = 3). **(B-C)** Venn diagrams display uniquely and shared rhythmic transcripts for WT and P2KO in active and quiescent conditions respectively. **(D)** Schematic representation of core clock machinery rhythmicity (traces for the individual transcripts provided in Supp. Fig. 7). **(E & H)** Polar plot representing the center-line peak of Active and Quiescent SGZ NSCs PER2-dependent mfuzz clusters (membership cutoff >0.5) of temporally co-expressed transcripts (names represent generalized biological processes based on overrepresentation GO analysis, full clustering and enrichment plots in Supp. Fig. 8 and Supp. Table 4). **(F-G)** Heatmaps ordered by phase (peak expression) displaying z-scored vst counts of Active WT and P2KO samples respectively. **(I-J)** Heatmaps ordered by phase (peak expression) displaying z-scored vst counts of Quiescent WT and P2KO samples respectively.

We applied the R package Mfuzz to identify temporally co-regulated transcripts (Supp. Fig. 8). Transcripts were assigned to 8 distinct clusters in both active and quiescent NSCs, based on manual inspection of different clustering numbers and their representations of existing patterns. Each cluster was subsequently analyzed for Gene Ontology enrichment (Count ≥ 5, FDR < 0.05, fold enrichment ≥ 2, minimum of 5 genes per pathway). While in Quiescent NSC clusters, the Mfuzz peak distribution is only bimodal, active NSCs clusters are more broadly distributed across circadian time (Fig 3. E & H, Supp. Fig. 8, Supp. Table 4).

Active NSC Clusters 2–5 peak between CT22 and CT2 and are enriched for glycogen and energy metabolism, lipid and organic acid metabolism, cell cycle (G2/M-mitotic), and neurogenesis genes. Cluster 6, enriched for ribosome biogenesis and rRNA processing genes, peaks at CT6, followed by Cluster 8 (CT14), enriched for RNA processing and cell cycle (G1/S-replication) regulation genes. Finally, Clusters 1 and 7 peak between CT15/CT16 and are enriched for cholesterol and sterol metabolism, ribosome biogenesis and tRNA processing genes (Fig. 3E, Supp. Fig. 8, Supp. Table 4). This temporal organization was exemplified by cell cycle regulators (Cluster 8: *Myc, Trp53, Ccnd1*) and neurogenic signaling components (Cluster 5: *Ntrk2, Epha4*), which showed ordered expression peaking in distinct, sequential time windows (Fig. 3F-G). With the exception of *Epha4* (2.37-fold change) expression levels of these markers were comparable between WT and P2KO samples, indicating that PER2 selectively governs their temporal organization rather than their absolute expression (Supp. Fig. 6D).

Quiescent PER2-dependent clusters display an enrichment peak mostly between CT0 and CT2 (CT: circadian time, i.e. constant conditions), highlighting a synchronization of PER2-dependent metabolic processes, with the exception of Cluster 2 (vacuolar/lysosomal organization), which peaked in antiphase at CT12 (Fig. 3 H). The clusters peaking between CT0 and CT2 are enriched for genes involved in sterol biosynthesis and cholesterol biosynthesis/metabolism (Clusters 3 & 6), ribosome biogenesis and RNA processing (Cluster 7), and organelle localization (Cluster 8). Clusters 1, 4 and 5 did not reach significance for any GO term (Count ≥ 5, FDR < 0.05, fold enrichment ≥ 2, minimum of 5 genes per pathway; Supp. Fig. 8, Supp. Table 4). This circadian organization was exemplified by lysosomal components (Cluster 2*: Lamp1*, *Creg1*, and *Tpp1*) peaking in antiphase to the coordinated expression of cholesterol and sterol synthesis components (Cluster 6: *Fdps*, *Dhcr7*, and *Scarb1*) alongside organelle localization and vesicle trafficking regulators (Cluster 8: *Ykt6*, *Stard3nl*, and *Uchl1*). This suggests a PER2-dependent temporal sequence of cholesterol synthesis, vesicle-mediated transport, and lysosomal activity in quiescent SGZ NSCs (Fig. 3Q, R). Again, *Per2* deletion abolished rhythmicity, but did not alter the expression levels of any of the aforementioned transcripts (Supp. Fig. 6 E).

While our primary focus was on PER2-dependent rhythmicity in WT NSCs we also applied clustering to transcripts uniquely rhythmic in P2KO but not rhythmic in WT samples (Supp. Fig. 9). Most of the quiescent rhythmic pathways overlapped with pathways rhythmic in WT quiescent NSCs (ncRNA processing, rRNA metabolism, and ribosome biogenesis) via different transcripts (Supp. Table 4). This suggests that the rhythmicity of these pathways is conserved by compensatory rhythmic mechanisms.

### 2.4 Circadian level changes of ASCL1 and CCND1 suggest coordinated cell cycle entry

Having identified key PER2-dependent rhythmic targets through our *in vitro* transcriptomic approach, we next sought to validate whether these circadian dynamics translate to the *in vivo* neurogenic niche. Among PER2-dependent rhythmic transcripts in active NSCs, we focused on Cyclin-D1 (CCND1) and Achaete-scute Family BHLH transcription factor 1 (ASCL1) for *in vivo* protein validation. CCND1 was selected as a downstream effector of BMAL1/CLOCK whose circadian regulation would predict phase-locked cell-cycle entry. ASCL1 was chosen as the primary transcriptional regulator of the quiescence-to-activation transition and neuronal fate commitment^30,31^. *Ccnd1* peaks with Cluster 8 and *Ascl1* peaks between different clusters of cell cycle transcripts (Clusters 4 and 8, Fig. 3 E).

In active NSCs, ASCL1 and CCND1 levels were elevated during daytime, peaking at ZT0, consistent with a clock-gated activation program (Figure 4A,B,E,F,G and J). In quiescent NSCs, ASCL1 and CCND1 protein levels were lower than in active cells, with no significant day-night variation (Fig. 4C,D,E,H,I and J). Together, these *in vivo* findings support a model in which key regulators of NSC activation and cell-cycle entry accumulate preferentially during the day, suggesting circadian priming for nighttime proliferation. We therefore next sought to evaluate cell-cycle entry as a functional readout of this temporal program and the potential involvement of PER2.

**Figure 4:**
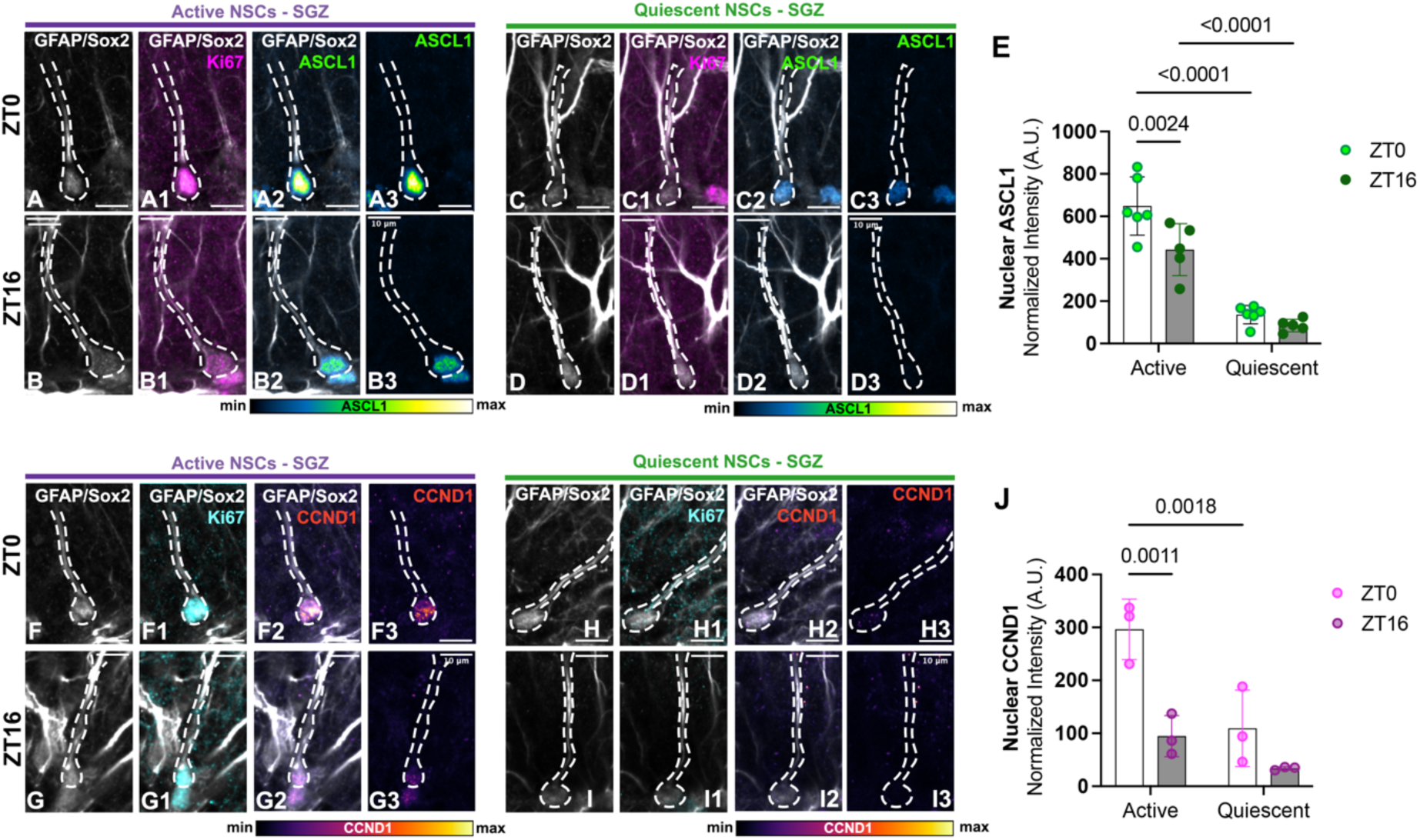
Daytime accumulation of ASCL1 and CCND1. (**A**-**D**) Active NSCs were identified as GFAP+ (single radial projection), nuclear Sox2+, proliferation marker **Ki67+**. Quiescent NSCs were identified as GFAP+, Sox2+, **Ki67-**. Nuclear masks were obtained and intensity measured in the channel of interest at both ZT0 (top panels) and ZT16 (lower panels). Dashed lines highlight NSCs in the SGZ. (**E**) Two-way ANOVA followed by Fisher’s LSD post-hoc test. Numbers represent p-value, dots display individual replicates and error bars represent SD. (**F-I**) Active and quiescent states were distinguished as previously described. (**J**) Two-way ANOVA followed by Fisher’s LSD post-hoc test. Numbers represent p-value, dots display individual replicates and error bars represent SD.

### 2.5 PER2 is required for circadian gated NSC proliferation in adult neurogenic niches

The PER2-dependent rhythmic pathways related to cell cycle regulation (*in vitro*), alongside the daytime-restricted expression of ASCL1 and CCND1 (*in vivo*), suggests circadian coordination of adult NSC activation and/or proliferation. Previous reports have demonstrated features of synchronized cell cycle entry in the SGZ ^15,17^ or SVZ^16^. Here, we analyzed this phenomenon from a comparative perspective to further elucidate niche-specific characteristics.

We first assessed whether adult NSC proliferation follows a diel pattern in both niches. Mice were treated with EdU (10 mg/ml, i.p.) 30 min before perfusion and sacrificed in 4h intervals from ZT0 to ZT20 (Fig. 5 A-B, E). Proliferation was assessed by EdU incorporation (S-phase cells) and Ki67 immunostaining (cell-cycle marker of late G1 to M phase^32^). The total number of Ki67+ cells in the SGZ remained unchanged across the day (Supp. Fig 10 B), whereas S-phase cells displayed a clear diel pattern, peaking between ZT16 and ZT20 (Fig. 5 C). In the SVZ, both Ki67+ (Supp. Fig. 10 C) and S-phase cell numbers showed a nighttime increase with the lowest levels of EdU+ cells between ZT0 and ZT4 and highest levels at ZT20 (Fig. 5 F).

**Figure 5:**
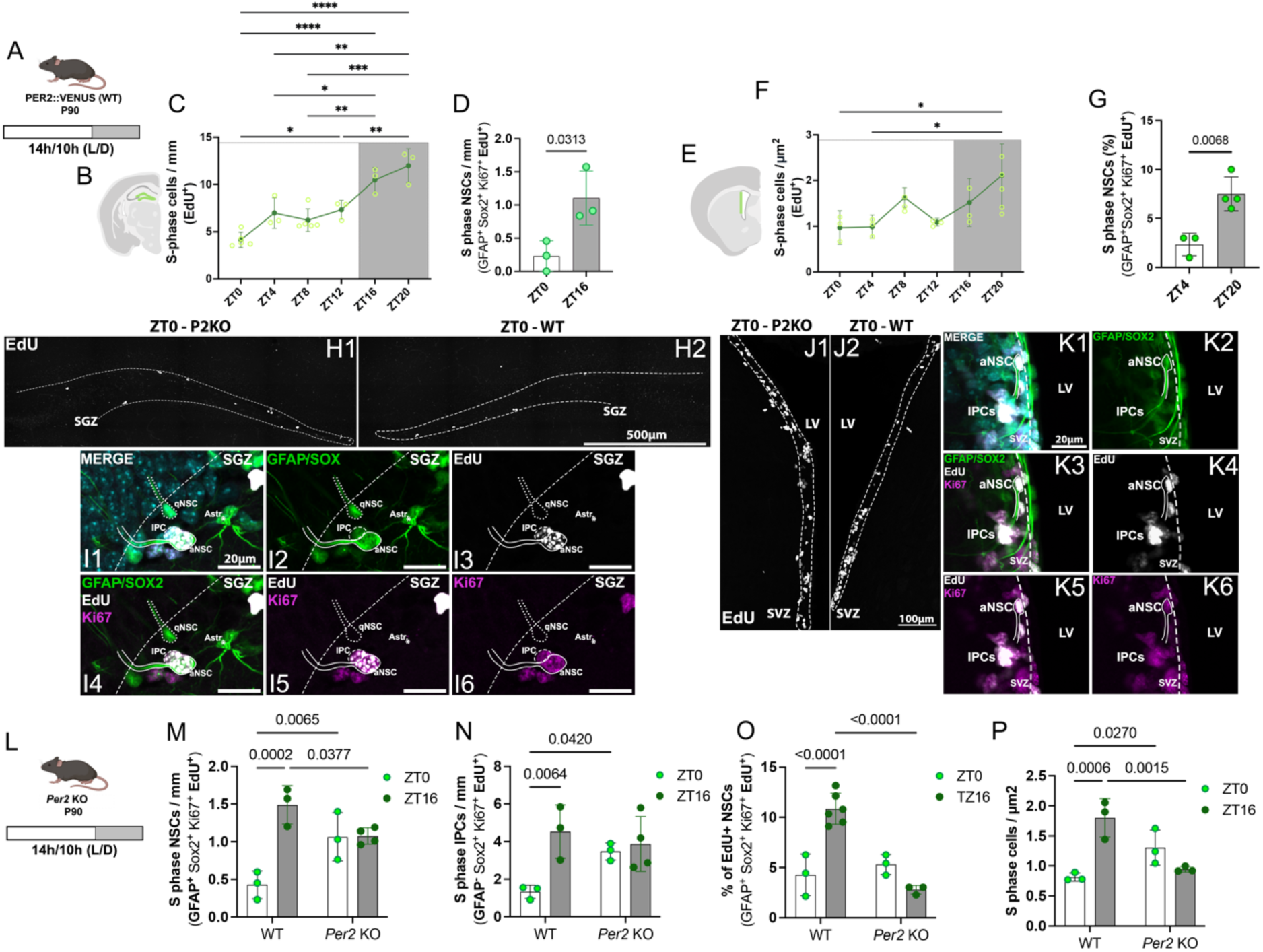
PER2-dependent coordination of NSC activation. **(A)** 90-day-old PER2::VENUS mice were kept in a 14 h/10 h light–dark cycle. **(B)** Green highlighted area represents ROI for SGZ quantifications. **(C)** Total number of EdU+ cells were significantly higher during night-time (one-way ANOVA followed by Tukey’s post hoc test, * p < 0.05, ** p < 0.005, *** p < 0.0005, **** p < 0.0001; brighter dots represent individual replicates, darker dots and bars represent average ± SD). **(D)** Number of S-phase NSCs (GFAP+ Sox2+ Ki67+ EdU+). **(E)** Green area highlights SVZ ROI for quantification. **(F)** Number of EdU+ cells were significantly higher at ZT20 (one-way ANOVA followed by Tukey’s post hoc test, * p < 0.05; brighter dots represent individual replicates, darker dots and bars represent average ± SD). **(G)** Percentage of S-phase NSCs (GFAP+ Sox2+ Ki67+ EdU+) was higher at ZT20 when compared to ZT4 (Unpaired t-test; values represent p values; dots represent replicates; error bars SD). **(H1–H2)** Immunostainings of EdU+ cells in the SGZ of WT and Per2 KO mice, respectively. Dashed line represents the limits of the SGZ within the dentate gyrus. **(I1–I6)** Identification of NSCs relied on location, morphology, and co-staining. **aNSC:** solid line highlights a S-phase NSC (single radial GFAP+ projection, Sox2+, Ki67+, EdU+). **IPC:** dashed line displays an S-phase IPC (GFAP−). **qNSC:** dotted line displays a quiescent NSC (GFAP+, Sox2+, Ki67−, EdU−). **(J1–J2)** Immunostainings of EdU+ cells in the SVZ of WT and Per2 KO mice, respectively. Dashed line represents the limits of the SVZ in the lateral ventricle (LV). **(K1–K6) aNSC:** S-phase NSCs were identified through a single GFAP+ radial projection and a Sox2+ nucleus (solid line; GFAP+, Sox2+, Ki67+, EdU+). **IPCs:** cluster of GFAP− Sox2+ Ki67+ EdU+ IPCs. **(L)** 90-day-old *Per2* KO mice were kept in a 14 h/10 h light–dark cycle. **(M)** Number of S-phase NSCs and is higher during night-time in wild-type animals and unchanged in *Per2* KO mice in the SGZ. **(N)** The same is observed for total the number of S-phase IPCs. (**O**) Percentage of S-phase NSCs in the SVZ displayed similar phenotype in WT animals but with reduced EdU+ NSCs during nighttime in P2KO animals. (**P**) Total number of S-phase cells was higher during nighttime in WT animals and constant in P2KO mice (two-way ANOVA followed by Fisher’s FSD post-hoc test, numbers represent p-values; dots display individual replicates; error bars SD).

To investigate whether coordinated cell-cycle entry was a feature of NSCs, we performed co-staining of these proliferation markers with GFAP and Sox2 in animals sampled during early daytime and nighttime. Active NSCs in the SGZ (ZT0 and ZT16) and SVZ (ZT4 and ZT20) were identified by the presence of a GFAP+ radial projection and their location in the niche (Fig. 5 I1-I6, K1-K6). In both SGZ and SVZ the number of S-phase NSCs was higher during nighttime (Fig. 5 D and G). Additionally, the number of EdU+ intermediate progenitor cells (IPCs) in the SGZ was higher during nighttime (Supp. Fig. 10 D).

We next utilized *Per2* KO animals to test whether the observed nighttime increase in S-phase NSCs is dependent on circadian clock functionality. In wild-type littermates, the number of S-phase NSCs and IPCs in the SGZ was significantly higher at ZT16 compared to ZT0; however, this difference was completely abolished in their *Per2* KO counterparts by an increase of proliferation during daytime (Fig. 5 M-N). In the SVZ, PER2-dependent effects were also observed. Like in the SGZ, the temporal variation in both the total number of S-phase cells (Fig. 5 P) and S-phase NSCs (Fig. 5 O) was lost in the absence of PER2, however, this loss of temporal gating was achieved by a decrease of proliferation during nighttime.

Taken together, our results reveal that diel patterns of NSC proliferation are present and dependent on PER2 in both the SGZ and SVZ niches, but through different mechanisms. Despite the absence of autonomous circadian clock synchronization in SVZ-derived cultures, *Per2* deletion abolished the diel organization of NSC proliferation in that niche. This implies that the underlying regulation is niche-specific.

## 3. Discussion

Together, our data demonstrate that PER2-dependent circadian mechanisms contribute to the temporal organization of NSC proliferation in the two main neurogenic niches of the adult mammalian brain. Our findings reveal different levels of circadian clock autonomy in SGZ and SVZ NSCs: circadian clock genes in SGZ-derived NSCs behave as self-sustained oscillators, whereas SVZ-derived NSCs lack detectable rhythmicity of circadian clock transcripts under the same conditions. The absence of detectable rhythmicity in cultured SVZ cells could indicate the absence of a cell-intrinsic oscillator. Alternatively, because cells are pooled in our assays, it can also reflect a lack of synchronization between those cells and/or low-amplitude rhythms.

We consider the latter possibility more plausible as we detect BMAL1 and PER2 oscillations in the NSCs of both niches *in vivo,* combined with observations from other adult stem-cell systems. Such a scenario is supported by *in vivo* data in mouse epidermal stem cells^8,33^. Consistently full physiological rhythmicity in the epidermis (including its stem cells) requires communication between central and tissue clocks^34^. Similarly, in enteroids, neighboring niche cells contribute to the coordination of stem-and progenitor-cell rhythms^35^. Thus, the absence of a detectable population-level rhythm does not necessarily indicate the absence of functional clocks in individual SVZ NSCs; rather, rhythmicity may persist at the single-cell level but remain undetectable in bulk *in vitro* measurements if cells are not synchronized.

In this context, identifying the cues that coordinate SVZ rhythmicity *in vivo*, and which may be absent from the *in vitro* environment, is an important next step. A previous report proposed melatonin as one such cue^16^. However, we observe robust clock-dependent circadian rhythms in melatonin-deficient mice (C57BL/6J background^18,19^), implying the existence of alternative signals. Glucocorticoid rhythms are also plausible candidates, given that corticosterone oscillations regulate NSC quiescence in the SGZ^36^, although their role in the SVZ remains to be further elucidated. Moreover, choroid plexus-derived factors regulate adult NSC activation^37^ and the choroid plexus translatome displays diurnal patterns associated with time-of-day-dependent changes in cerebrospinal fluid (CSF) composition^38^. Together, these observations identify melatonin, glucocorticoids, and CSF-derived factors as potential contributors to the temporal coordination of the SVZ niche. Further investigation is needed to determine how systemic cues, local niche-derived signals, and central circadian inputs interact to synchronize rhythmicity within each neurogenic niche and how resilient NSC clocks are to the absence of such cues.

Our data reveal cell-state-dependent differences in the temporal dynamics of core circadian clock components. *In vivo*, both BMAL1 and PER2 displayed rhythmic protein dynamics in total and quiescent NSCs, whereas active NSCs displayed predominantly PER2 rhythmicity, with BMAL1 levels remaining relatively low and not differing between time points in either niche. Consistently, SGZ-derived cells exhibited state-specific biases: active NSCs retained rhythmicity in negative-arm components, including PER2::VENUS and *Per2*, *Cry2*, and *Bhlhe41* transcripts, whereas quiescent NSCs displayed rhythmicity in positive-arm components, including BMAL1 protein and *Clock* and *Bmal1* transcripts. While further analysis will be needed, this pattern suggests state-dependent differences in the synchrony and/or amplitude of distinct clock arms within active and quiescent NSC populations, with potential implications for the temporal regulation of NSC state transitions and maintenance.

Our *in vitro* system allowed us to investigate NSCs in the absence of ongoing systemic and central circadian inputs. The persistence of transcriptomic circadian rhythms under these conditions shows they are maintained in cultured NSCs, either through cell-intrinsic mechanisms or through local interactions retained *in vitro*. Comparisons between WT and *Per2*-KO (P2KO) NSCs identified PER2-dependent rhythmic transcription in quiescent and active SGZ-derived NSCs. In quiescent NSCs, PER2-dependent rhythmic transcripts mainly belong to cholesterol metabolism and lysosomal function (well-established quiescence pathways^39–41^) and organelle localization, indicating it as a novel quiescence feature. Future work will determine how circadian time and phase relationships can regulate the state of quiescence. In active SGZ-derived NSCs, PER2-dependent rhythmicity converged on cell-cycle control (*Trp53*, *Ccnd1*, *Myc*) and the proneural regulator *Ascl1*, pointing to a clock output directed at proliferative timing in active NSCs. It is unlikely that the observed circadian oscillations are a product of the characterized *in vitro* ultradian rhythms (∼3 h), as these are not synchronized^31^. The asynchrony of the ultradian rhythms provide heterogeneity to NSC activation despite an overall increase in ASCL1 levels driven by the circadian component.

*Per2* deletion disrupted the temporal distribution of S-phase NSCs in both neurogenic niches, although through distinct mechanisms: increasing daytime proliferation in the SGZ while reducing nighttime proliferation in the SVZ. Thus, although PER2 is required for diel coordination of proliferation in both niches, its functional consequences appear to be shaped by niche-specific regulatory mechanisms. Whether the altered proliferation observed following *Per2* deletion reflects disrupted rhythmicity in effectors such as ASCL1 and CCND1 remains to be determined.

As a final point, we wondered, how rhythmic transcripts and pathways can persist in SVZ NSCs *in* vitro despite no detectable rhythmicity in core circadian clock components. As these rhythms were detected at the population level, they indicate that at least a subset of SVZ-derived cells retained sufficient temporal coordination to generate coherent transcriptional oscillations. One possibility is that rhythmic outputs are driven by low-amplitude or incompletely synchronized core circadian clock oscillations that fall below the detection threshold of our assays. Alternatively, they may arise through mechanisms that do not require detectable oscillations of canonical clock components, including metabolic or other TTFL-independent oscillatory processes^42–45^. Future work should define the mechanisms underlying these rhythmic SVZ outputs, determine whether comparable core-clock-independent rhythms are also present in SGZ-derived NSCs, and establish whether such mechanisms contribute to regional differences in circadian transcriptional organization.

### Limitations of the study

Our *in vitro* analyses were performed at the bulk-population level and therefore could not distinguish a loss of circadian clock rhythmicity from reduced synchrony among individual cells, which could be further elucidated through live-cell imaging and single-cell tracking. Our functional analyses focused primarily on the temporal coordination of cell-cycle progression, an important regulator of adult NSC activity. However, the rhythmic pathways identified in this study likely influence additional aspects of NSC biology that were not directly examined. In turn, how these aspects of NSC biology (e.g., fate decisions) impact circadian rhythms themselves remains to be determined.

A further limitation is that, although the characterization of PER2 and BMAL1 dynamics supports circadian clock function *in vivo*, other core clock components and auxiliary feedback loops were not assessed. Similarly, while *Per2* knockout enabled the identification of PER2-dependent phenotypes and rhythmic transcriptional programs, the contributions of other core circadian clock components were not examined and likely differentially regulate NSCs. Finally, although *Per2* deletion clearly disrupted NSC dynamics, its long-term effects on NSC pool maintenance were not assessed. This will be particularly important to determine in a niche-specific manner, given the divergent proliferation phenotypes observed in the SGZ and SVZ and the potential relevance of circadian disruption to adult neurogenesis and human health.

## Acknowledgements

We would like to acknowledge Vinoth Babu Veedin Rajan for contributions in the initial phase of the project and to thank all the Urbán Lab members who supported sampling efforts, Amarbayasgalan Davaatseren, Lidija Milojković, Justus Kleifeld, Greeshma Pushpa Bose, and sampling processing, Juliane Baar. We would like to extend our appreciation for the support of the BioOptics, NGS and Histology facilities at the VBCF, Histology facility at Max Perutz Labs and the staff of the animal facility IMBA/IMP.

## Contributions

**Pedro O. Brum:** Conceptualization; data curation; formal analysis; validation; investigation; visualization; methodology; writing -original draft; writing - review & editing.

**Ava Abdi:** Data curation; formal analysis; validation; investigation; visualization; methodology; writing - review & editing.

**Federico Scaramuzza:** Data curation; formal analysis; writing - review & editing.

**Tatjana Kepcija:** Methodology; resources; writing - review & editing.

**Michael Hastings:** Resources; writing – review & editing

**Andrew Loudon:** Resources; writing – review & editing

**Kristin Tessmar-Raible^#^:** Conceptualization; resources; formal analysis; supervision; funding acquisition; investigation; project administration; writing - original draft; writing - review & editing.

**Noelia Urbán^#^:** Conceptualization; resources; formal analysis; supervision; funding acquisition; investigation; project administration; writing - original draft; writing - review & editing.

#The order of the corresponding authors (KT-R and NU) was decided through a game of heads or tails.

---

**Supplemental Figure 1:**
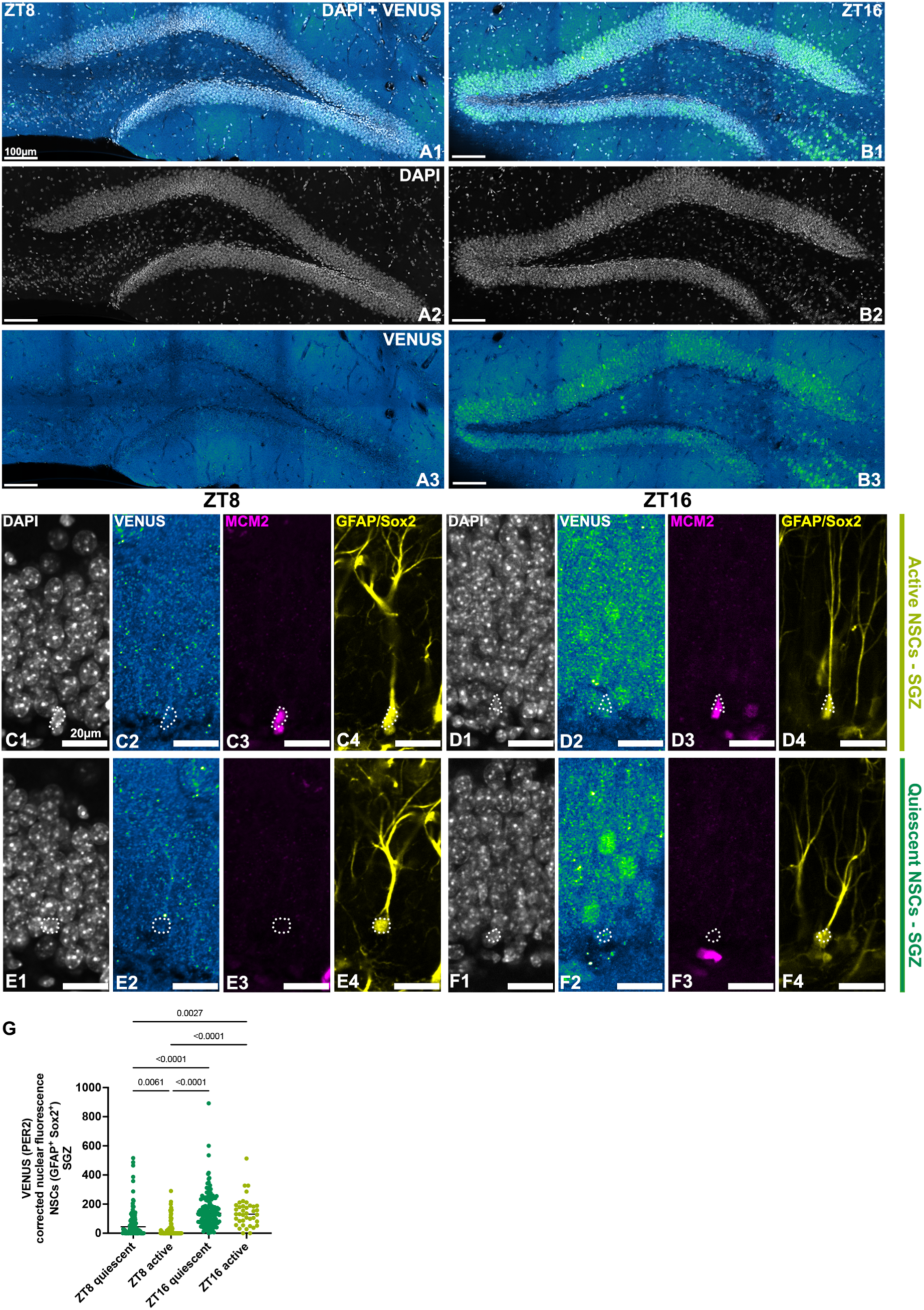
Dentate Gyrus **(A1-A3)** VENUS (PER2) staining at ZT8 and **(B1-B3)** and ZT16 highlighting the general trend of elevated levels of PER2 at nighttime, including cells in the granule layer, hilus and SGZ. Active NSCs (GFAP+ Sox2+ MCM2+) VENUS (PER2) expression at **(C1-C4)** ZT8 and **(D1-D4)** ZT16. Quiescent NSCs (GFAP+ Sox2+ MCM2-) VENUS (PER2) expression at **(E1-E4)** ZT8 and **(F1-F4). (G)** Dot plot of background corrected nuclear fluorescence of quiescent and active NSCs at ZT8 and ZT16 (N=4-5 animals per group). Welch ANOVA followed by Games-Howell multiple comparison test. Values represent p value.

**Supplemental Figure 2:**
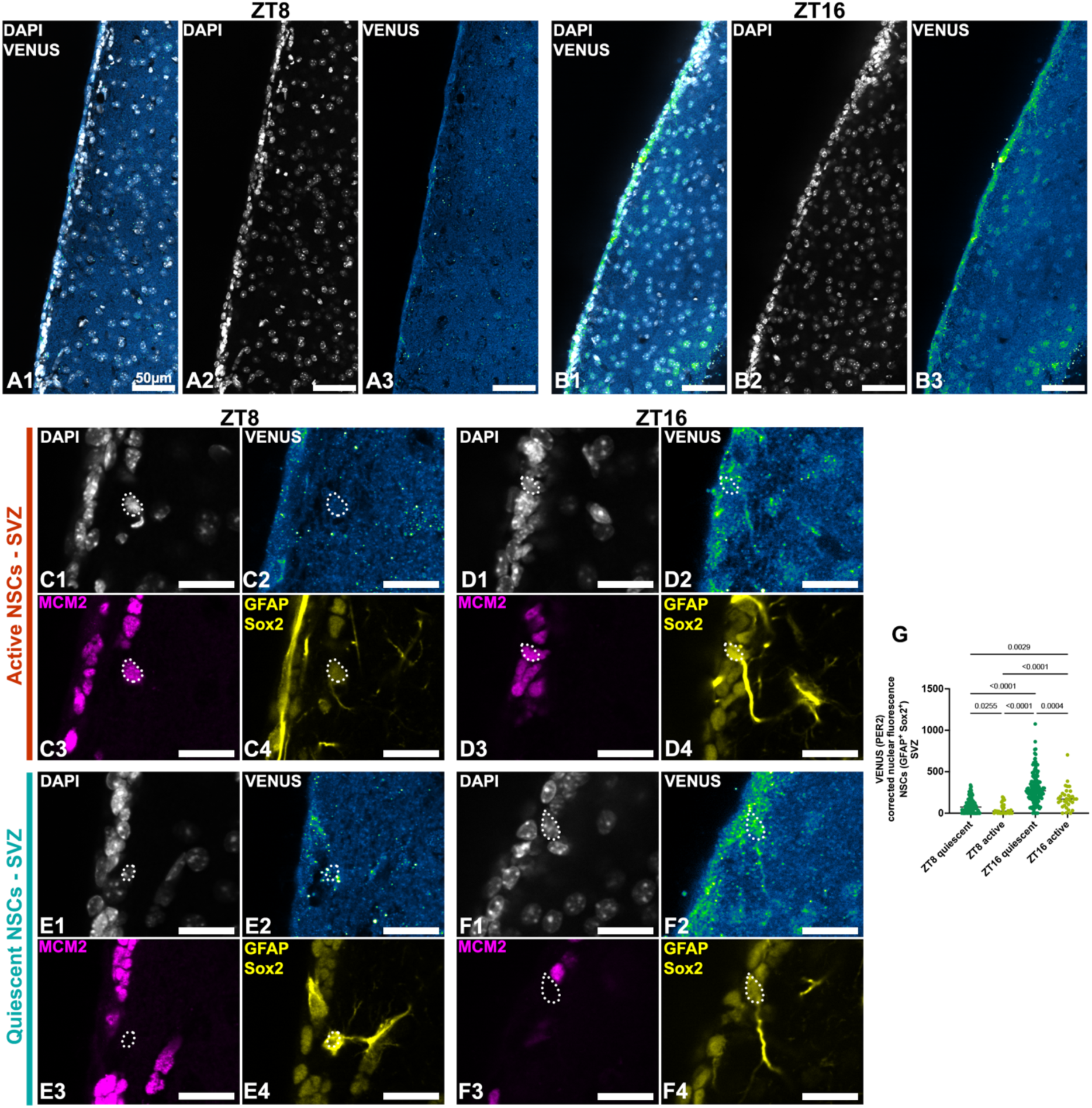
Lateral Ventricle **(A1-A3)** VENUS (PER2) staining at ZT8 and **(B1-B3)** and ZT16 highlighting the general trend of elevated levels of PER2 at nighttime including cells along the ventricle wall and the cells adjacent to the SVZ. Active NSCs (GFAP+ Sox2+ MCM2+) VENUS (PER2) expression at **(C1-C4)** ZT8 and **(D1-D4)** ZT16. Quiescent NSCs (GFAP+ Sox2+ MCM2-) VENUS (PER2) expression at **(E1-E4)** ZT8 and **(F1-F4). (G)** Dot plot of background corrected nuclear fluorescence of quiescent and active NSCs at ZT8 and ZT16 (N=3-4 animals per group). Welch ANOVA followed by Games-Howell multiple comparison test. Values represent p value.

**Supplemental Figure 3:**
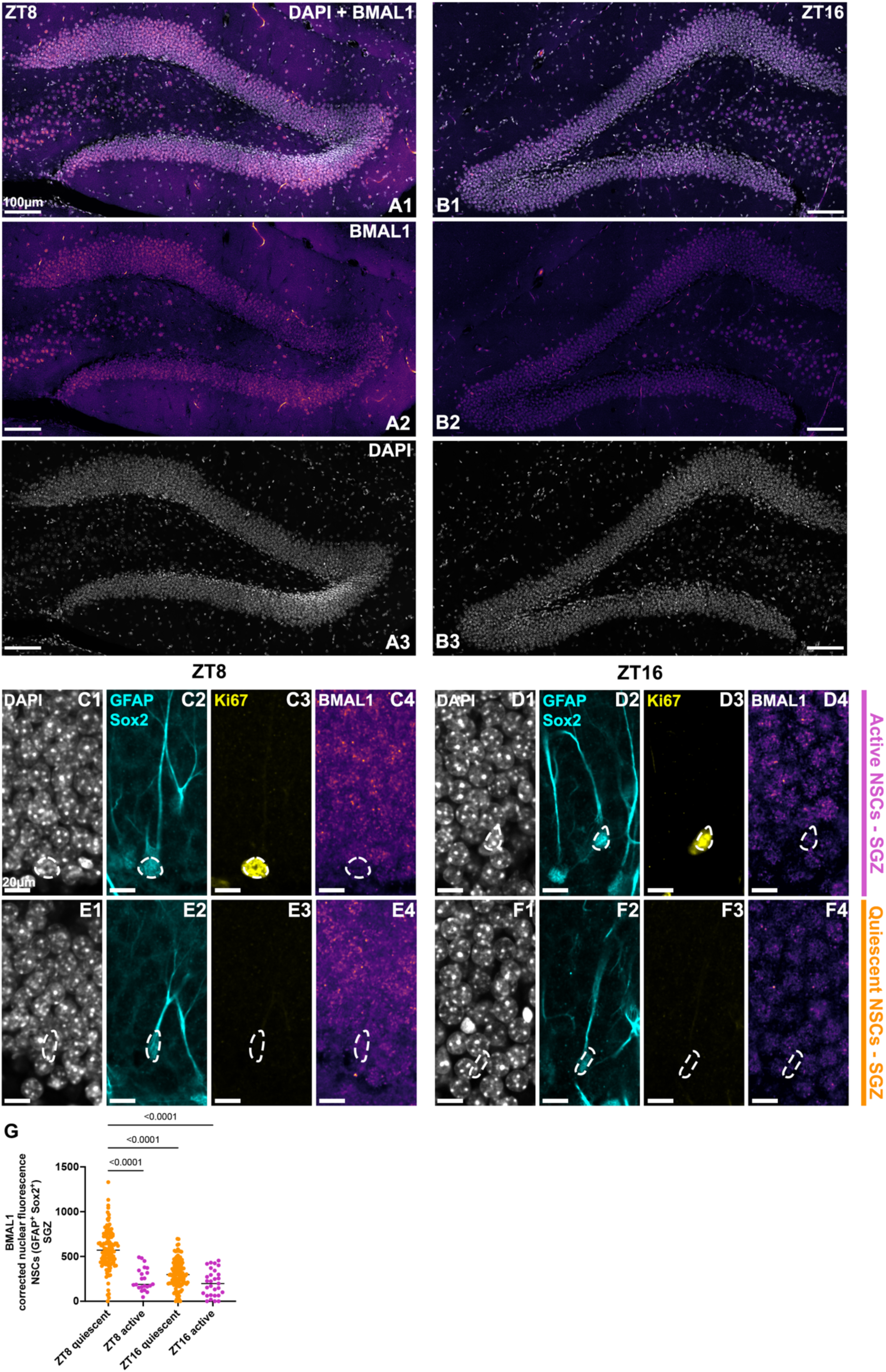
Dentate Gyrus **(A1-A3)** BMAL1 staining at ZT8 and **(B1-B3)** and ZT16 highlighting the general trend of elevated levels of BMAL1 during daytime including cells in the granule layer, hilus and SGZ. Active NSCs (GFAP+ Sox2+ Ki67+) BMAL1 expression at **(C1-C4)** ZT8 and **(D1-D4)** ZT16. Quiescent NSCs (GFAP+ Sox2+ Ki67-) BMAL1 expression at **(E1-E4)** ZT8 and **(F1-F4)** ZT16**. (G)** Dot plot of background corrected nuclear fluorescence of quiescent and active NSCs at ZT8 and ZT16 (N=4 animals per group). Welch ANOVA followed by Games-Howell multiple comparison test. Values represent p value.

**Supplemental Figure 4:**
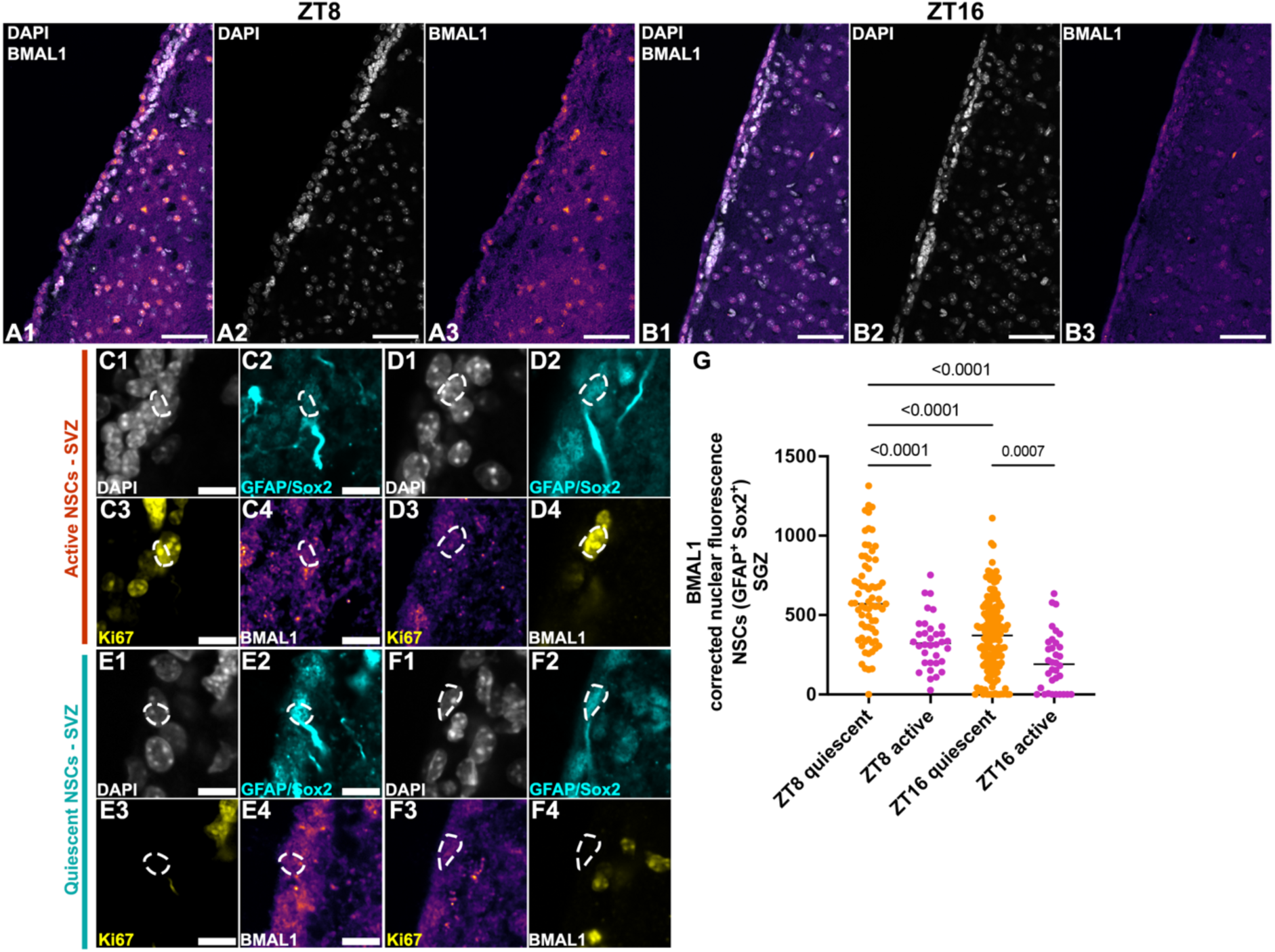
SVZ **(A1-A3)** BMAL1 staining at ZT8 and **(B1-B3)** and ZT16 highlighting the general trend of elevated levels of BMAL1 during daytime including cells in the granule layer, hilus and SGZ. Active NSCs (GFAP+ Sox2+ Ki67+) BMAL1 expression at **(C1-C4)** ZT8 and **(D1-D4)** ZT16. Quiescent NSCs (GFAP+ Sox2+ Ki67-) BMAL1 expression at **(E1-E4)** ZT8 and **(F1-F4)** ZT16**. (G)** Dot plot of background corrected nuclear fluorescence of quiescent and active NSCs at ZT8 and ZT16 (N=4 animals per group). Welch ANOVA followed by Games-Howell multiple comparison test. Values represent p value.

**Supplemental Figure 5:**
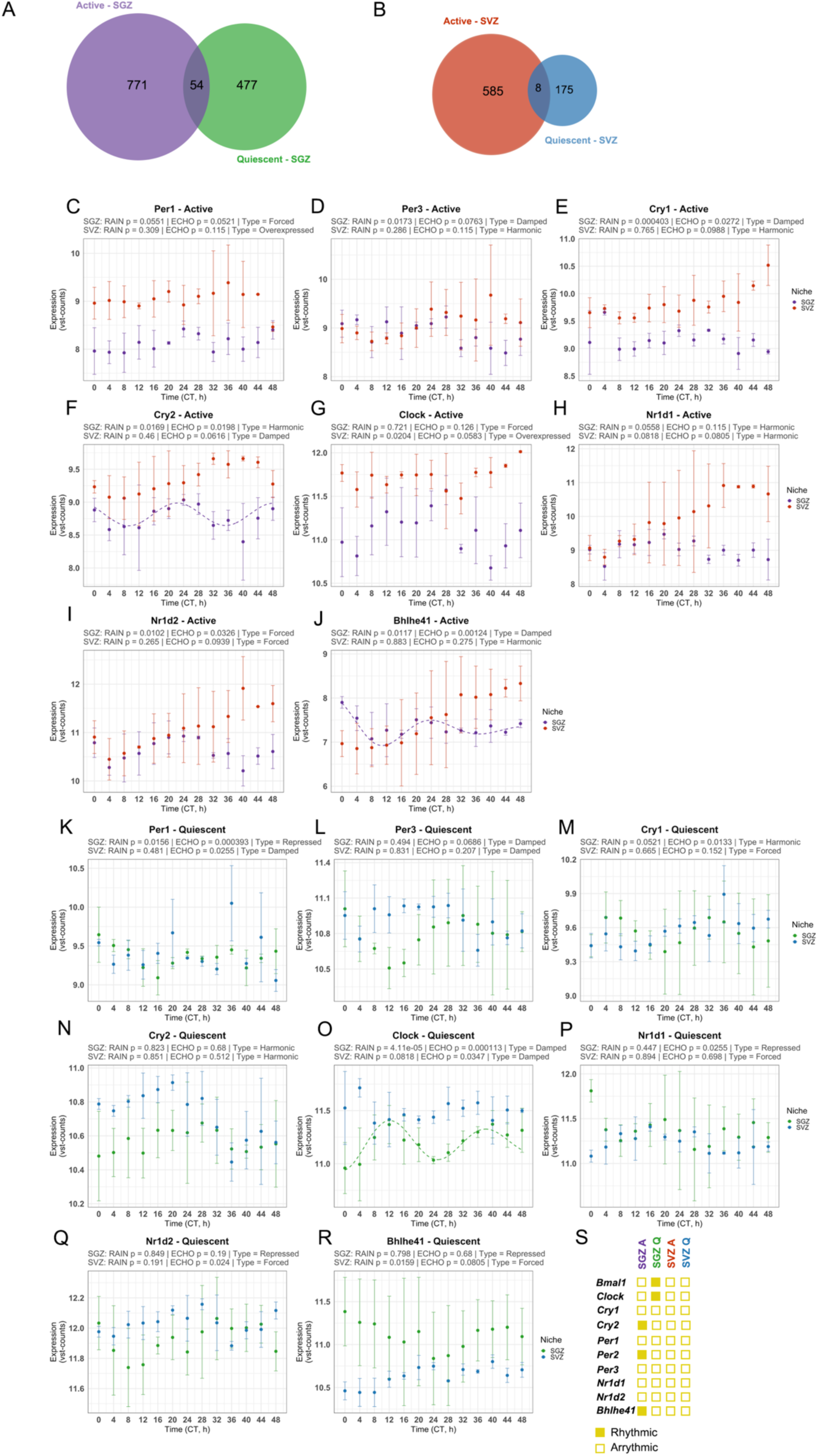
Venn diagrams display uniquely and shared rhythmic transcripts in Active and Quiescent of both (**A**) SGZ and (**B**) SVZ. **(C-J)** Core clock plots for active NSCs and **(K-R)** quiescent NSCs.

**Supplemental Figure 6:**
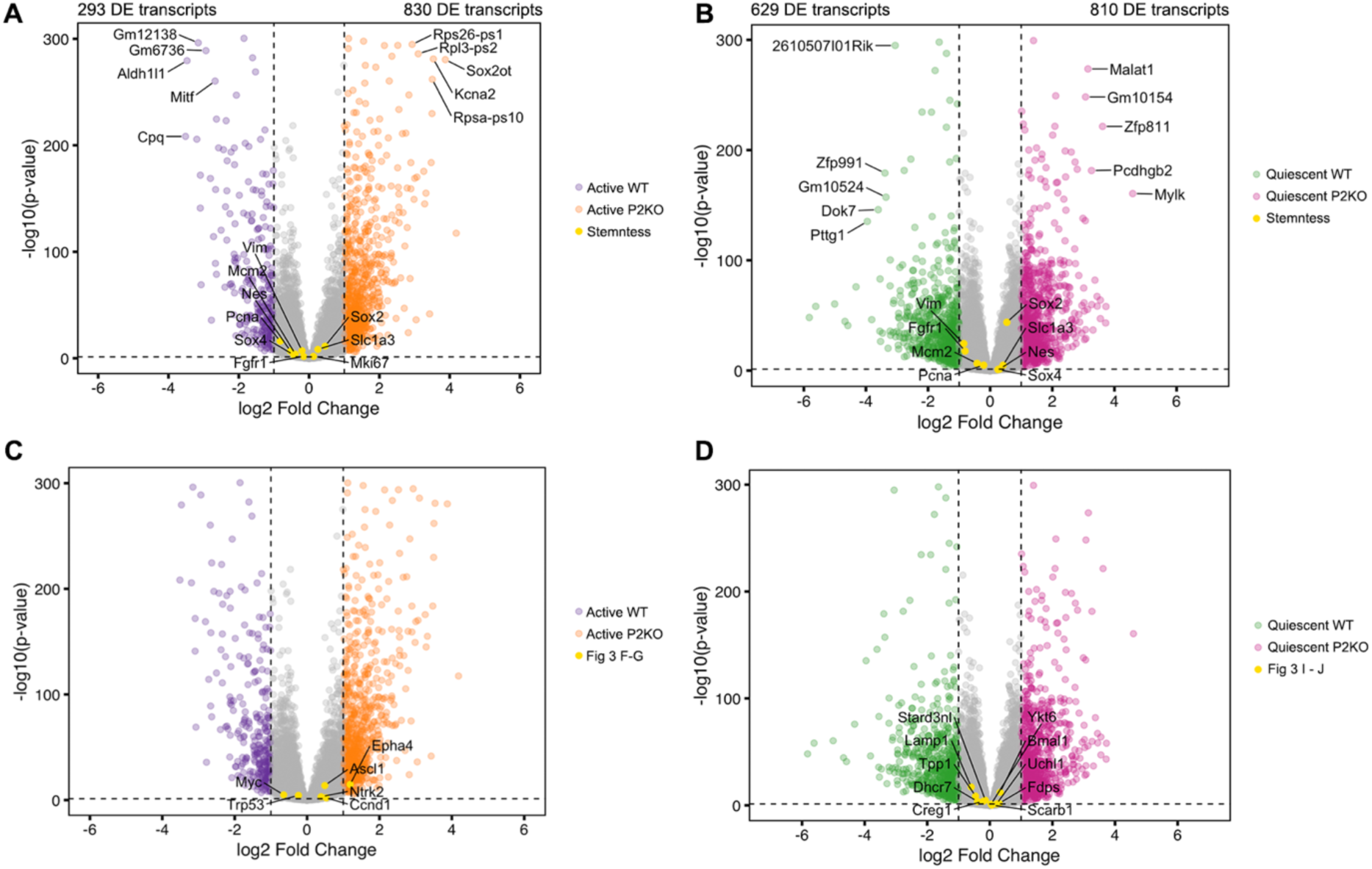
(**A**) Volcano plot showing differentially expressed (DE) transcripts between active WT and active P2KO neural stem cells. (**B**) Volcano plot showing DE transcripts between quiescent WT and quiescent P2KO neural stem cells. Yellow dots indicate stemness markers. The top five DE transcripts are annotated in each volcano plot. (**C**) Same volcano plot as in A, highlighting genes associated with the comparison shown in **Fig. 3F–G**. (**D**) Same volcano plot as in B, highlighting genes associated with the comparison shown in **Fig. 3I–J**. Dashed lines indicate the thresholds used for statistical significance and fold change (FDR < 0.05 and |log2 fold-change| ≥ 1).

**Supplemental Figure 7:**
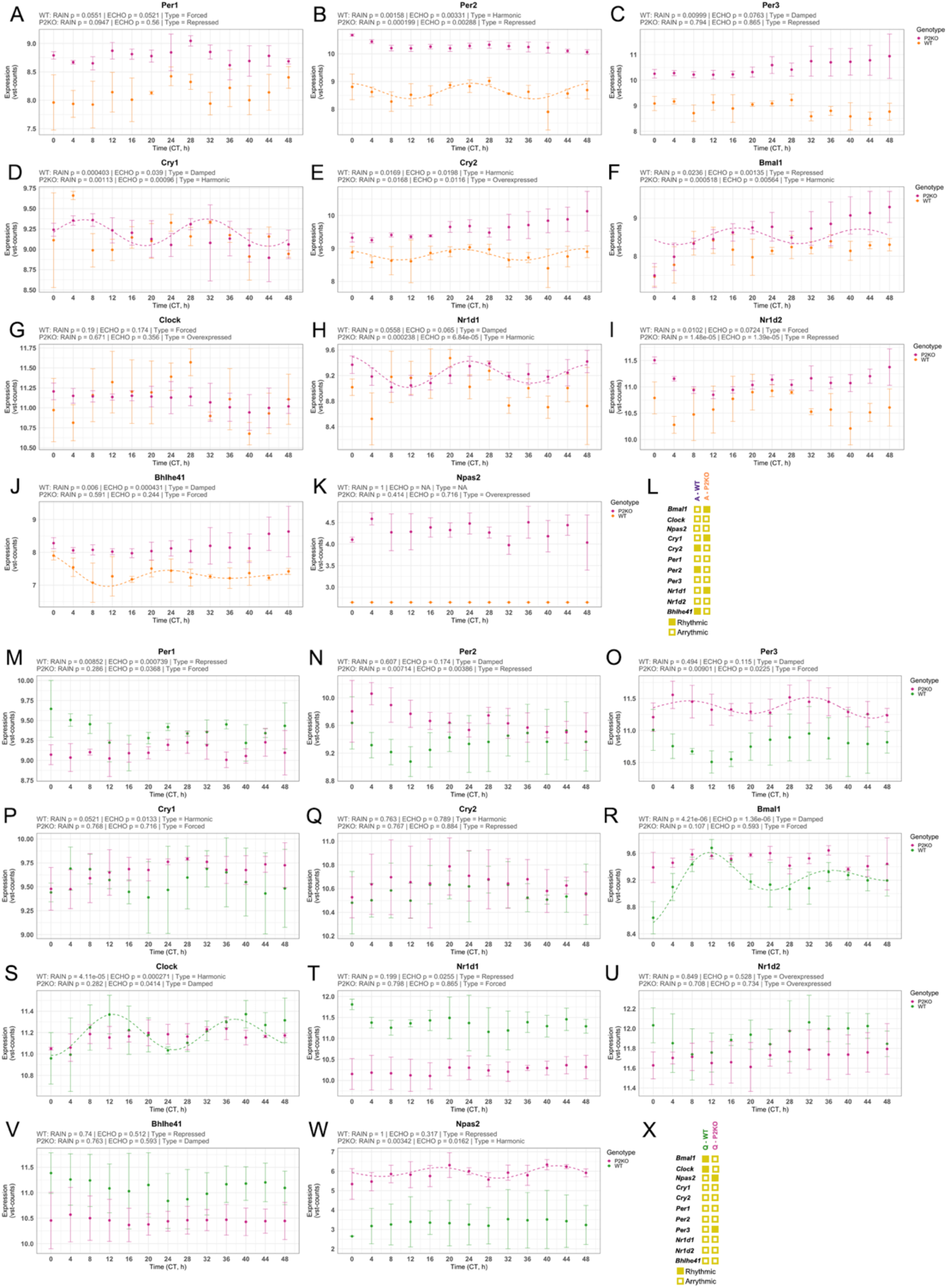
Core clock plots for **(A-K)** active SGZ NSCs and **(M-W)** quiescent SGZ NSCs respective. (**L** & **X**) Schemes displaying rhythmic and arrhythmic transcripts for active and quiescent NSCs.

**Supplemental Figure 8:**
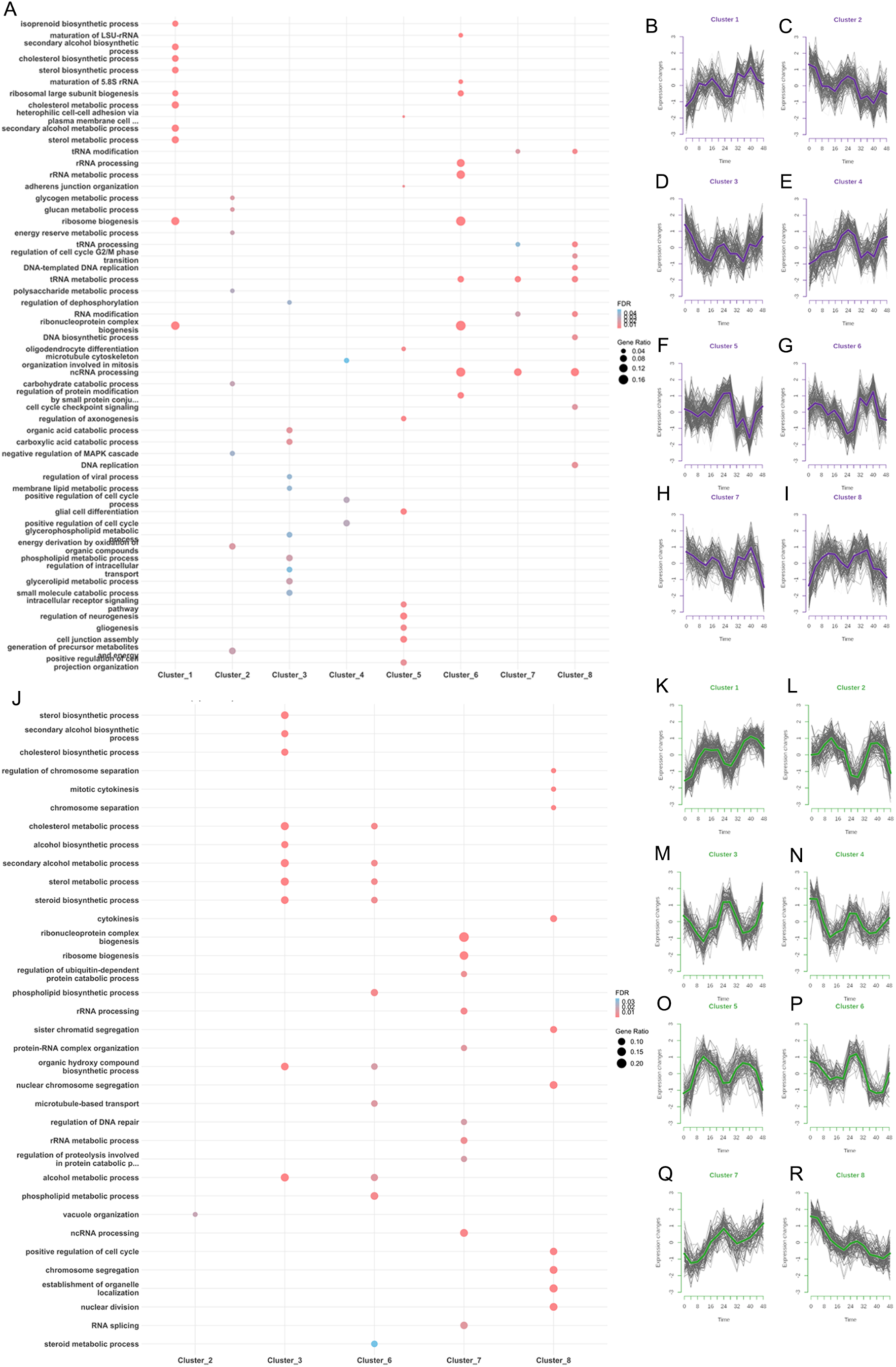
**(A)** Top 10 enriched pathways for all active SGZ clusters. **(B-I)** Mfuzz clusters (membership cutoff > 0.5) of PER2-dependent transcripts in active SGZ-derived NSCs. **(J)** Top 10 enriched pathways for all quiescent SGZ clusters. **(K-R)** Mfuzz clusters of PER2-dependent transcripts in quiescent SGZ-derived NSCs. Full list of enrichment in Supp. Table 4.

**Supplemental Figure 9:**
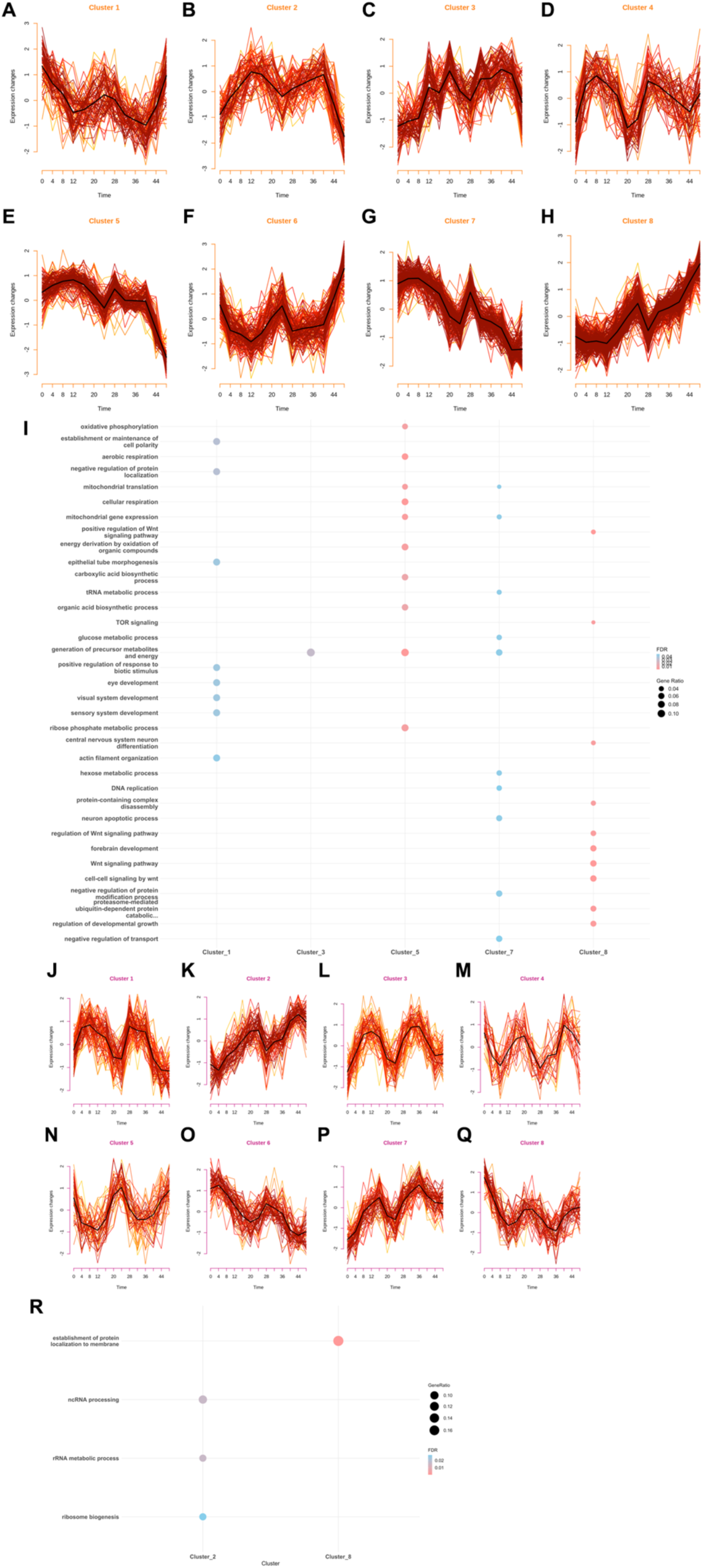
**(A-H)** Mfuzz clusters (membership cutoff > 0.5) of P2KO transcripts in active SGZ-derived NSCs. **(I)** Top 10 enriched pathways for all active SGZ clusters. **(J-Q)** Mfuzz clusters of PER2-dependent transcripts in quiescent SGZ-derived NSCs. Full list of enrichment in Supp. Table 4. **(R)** Top 10 enriched pathways for all quiescent SGZ clusters.

**Supplemental Figure 10:**
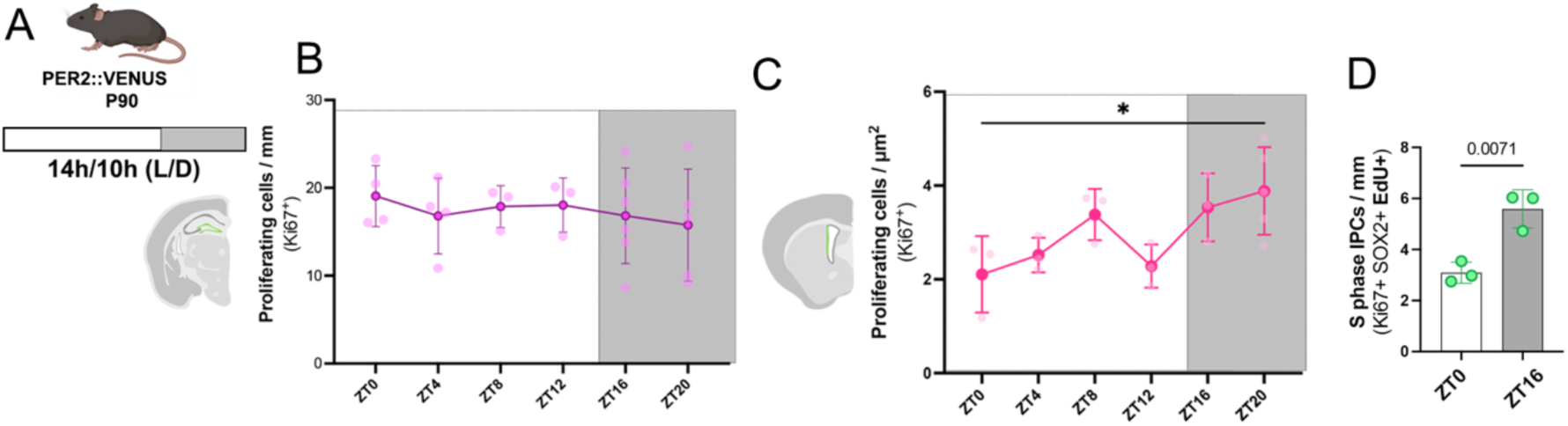
**(A)** P90 PER2::VENUS mice were kept in a 14h/10h (L/D). Green area displays ROI for quantifications in the SGZ. **(B)** Total proliferating cells in the SGZ normalized by the length of the SGZ. **(C)** ROI for quantifications of the SVZ and total number of proliferating cells normalized by the area (One-way ANOVA followed by Tukey’s *post hoc* test; * *p* < 0.05). **(D)** S-phase IPCs were increased during nighttime (Unpaired t-test; value represent p-value; dots represent replicates; error bars SD).

## 4. Methods

### 4.1 Animal husbandry and mouse models

All animals were housed and bred in accordance with Austrian and European legislation under approved ethical animal license protocols. PER2::VENUS mice were imported as embryos from the MRC Laboratory of Molecular Biology, Cambridge (provided by Michael Hastings, MRC; co-developed with Andrew Loudon, University of Manchester^21^). Per2 knockout (P2KO) mice were obtained from Jackson Laboratories (RRID:IMSR_JAX:010832). C57BL/6J mice were sourced from the internal colony at the Comparative Medicine Facility (IMBA/IMP, Vienna). All strains were maintained under a 14:10 h light/dark cycle with food and water available *ad libitum*, housed in individually ventilated cages (Tecniplast GM500) with a maximum of 5 animals per cage depending on body weight, at a constant temperature of 24°C.

### 4.2 Tissue collection and processing

Animals received a single intraperitoneal injection of EdU (10 mg/ml; Carl Roth, cat.: 7845.6) 30 min prior to euthanasia. Under anesthesia, animals were transcardially perfused with phosphate-buffered saline (1X PBS) until complete clearance of blood, followed by 10 min perfusion with 4% paraformaldehyde (PFA; Sigma, cat. no. P5148). Perfusions were performed every 4 h from ZT0 (defined as the moments lights were turned on) to ZT20. For night timepoints (ZT16 and ZT20), red-light LEDs were used to minimize the impacts of light exposure during tissue collection. Brains were washed in 1X PBS and post-fixed in 4% PFA overnight at 4°C with gentle agitation. Coronal sections (40 µm) were obtained using a vibratome (Leica VT1000S) and stored free-floating in 1X PBS with 0.02% Sodium Azide (ITW Reagents, A1430).

**For immunostainings,** brain sections were blocked for 1h 30min at room temperature with 1% Triton-PBS with 10% normal goat serum (NGS, G9023). Primary and secondary antibodies were diluted in solutions with the same composition. Primary antibody incubation was performed overnight at 4°C and followed by 5 washes of 10 minutes with 0.1% Triton-PBS. Secondary incubation was performed at room temperature for 2h and followed by additional 5 washes with 0.1% Triton-PBS. Finally, samples were incubated with 1 μg/ml DAPI (Sigma, D9542) in 1:1 PBS:H_2_O at room temperature for 30 min. Sections were mounted using AquaPolymount (Polysciences, 07918606-5) and let dry overnight. EdU detection was performed before primary incubation when necessary, following manufacturer’s instructions (Invitrogen, C10641). Samples were imaged utilizing an Olympus spinning disc confocal microscope (Olympus IX83 inverted microscope) equipped with a Yokogawa W1 and a Hamamatsu ORCA Fusion Digital CMOS camera (Hamamatsu Photonics K.K., C14440-20UP). Z-stacks and MIA were acquired with a 40x/0.75 air objective (Olympus) covering 30 μm of the specimen with 1-μm intervals. Images were stitched live with CellSens using automated panoramic imaging (auto MIA).

### 4.3 Image quantifications

***Neural Stem Cell (NSC) Identification*** (Fig. 1, 4, 5, Supp. Fig. 1-4, 10) preceded EdU/Ki67 co-staining analysis and nuclear intensity analysis and it was performed as following:

***Subgranular Zone (SGZ):*** Neural stem cells (NSCs) in the SGZ were identified by the presence of a single radial GFAP⁺ projection, a Sox2⁺ nucleus, and localization beneath the granule cell layer of the dentate gyrus (DG). To avoid ambiguous classification of astrocytes, cells in which the radial projection bifurcated within a range of two nuclear diameters toward the granule cell layer were excluded from NSC quantification^25^. Quantification in the SGZ were normalized by the length of the area of interest. Length was calculated by manually outlining the SGZ using the segmented line tool in Fiji^46^.

***Subventricular Zone (SVZ):*** Given the inherent challenges of NSC identification in coronal sections of the SVZ, cells were classified as NSCs based on their characteristic radial morphology and co-expression of GFAP+ and Sox2+^23,24^. As this approach does not capture the total NSC population, cells in which morphology was not clearly discernible due to sectioning orientation were excluded from quantification, values are reported as proportions of the total analyzed population rather than absolute counts. To improve reproducibility and focus on the neurogenic lineage, analysis was restricted to the upper 1,000 µm of the lateral wall.

***S-phase NSCs*** were defined by the criteria aforementioned in combination with EdU incorporation. ***Active NSCs*** were defined as Ki67⁺. ***Quiescent NSCs*** were defined as Ki67-. Intermediate progenitor cells (***IPCs***) were identified as Sox2⁺/Ki67⁺ cells lacking radial projections, and ***S-phase IPCs*** as the EdU⁺ subset of this population.

***For intensity quantifications (Figs. 1* *and* *4*)**, the z-plane of highest DAPI intensity was selected for each nucleus. Nuclei were manually segmented and mean gray values were measured. Nuclear segmentation was performed based on DAPI, GFAP, and Sox2 channels to confirm cell identity, with the investigator blinded to the channels of interest (PER2::VENUS and BMAL1, ASCL1 and CCND1) during ROI definition. Background subtraction was performed by measuring fluorescence intensity in the nearest cell-free region using a mask of identical area to the nuclear ROI, which was then subtracted from the nuclear intensity value. **Fig. 1**: 30-40 NSCs per animal across 3-4 slices (10 NSCs/slice) were randomly selected for quantification. **Fig. 4**: All the active NSCs across 6 slices were segmented for quantification and a matched number of randomly selected quiescent NSCs.

***Total EdU and Ki67*** (Fig. 5, Supp. Fig. 10):

**SGZ:** All EdU+ cells in the SGZ were quantified and normalized by the length of the area of interest.

**SVZ:** The upper 1000 μm of the lateral wall was included in the quantification, and the numbers were normalized to the SVZ area. The area was acquired by manually outlining solely the portion of SVZ being quantified in the maximum-intensity projection of hyperstacks using the segmented line tool in Fiji^46^.

### 4.4 Statistical analysis – *in vivo* datasets

Statistical analyses were performed using Prism (v10.0; GraphPad Software Inc., San Diego, USA). For time-series data comprising all 6 sampled timepoints, a one-way ANOVA with Tukey’s post-hoc multiple comparisons test was applied. Comparisons between two groups were performed using unpaired t-tests, and two-way ANOVA followed by Fisher’s FSD post-hoc test was used when comparing timepoints across genotypes (WT vs P2KO) and conditions (Active vs Condition).

### 4.5 Cell culture

Primary adult NSC cultures were established from 7-week-old mice. After cervical dislocation and brain extraction, we dissected both lateral ventricle^23^ and dentate gyrus^47^, and dissociated the tissue using the Neural Tissue Dissociation Kit (P) (Miltenyi Biotec, cat. no. 130-092-628). NSCs were initially expanded under non-adherent conditions as neurospheres in basal medium (DMEM/F-12, GlutaMAX™ supplement; Thermo Fisher Scientific, cat. no. 11559726) supplemented with 1× NeuroCult Supplement (STEMCELL Technologies, cat. no. 05701), 1× Penicillin/Streptomycin (Thermo Fisher Scientific, cat. no. 15140), 20 ng/ml FGF2 (EUBIO, cat. no. 450-33), 20 ng/ml EGF (PeproTech, cat. no. 315-09), and 2 µg/ml Heparin (Sigma-Aldrich, cat. no. H3393) for a minimum of 2 passages. Neurospheres were subsequently dissociated and transferred to 2D adherent cultures. Adherent NSCs were maintained in a proliferative state in basal medium consisting of DMEM/F-12, GlutaMAX™ supplement (Thermo Fisher Scientific, cat. no. 11559726), 20 ng/ml FGF2 (EUBIO, cat. no. 450-33), 20 ng/ml EGF (EUBIO, cat. no. 315-09), N2.1 supplement (Molecular Biology Service, VBC), 1× Penicillin/Streptomycin (Sigma-Aldrich, cat. no. P4333), 5 µg/ml Heparin sodium salt (Sigma-Aldrich, cat. no. H3393), and 2 µg/ml Laminin (Sigma-Aldrich, cat. no. L2020). All cultures were maintained at 37°C in 5% CO2.

### 4.6 Sampling for RNA sequencing and Immunostainings

On D0 (Fig. 2 A, Fig. 3 A), cells were plated in T25 flasks at 30.000 cells/cm² (RNA sequencing) or glass coverslips (Immunostainings, VWR, cat.: 631-0149) pre-coated with laminin (2μg/ml, Sigma-Aldrich, L2020, 15min/flasks, 2h/coverslips) in maintenance culture conditions (FGF2 + EGF as described above). After 24h, media was changed to quiescence induction media: DMEM/F-12, GlutaMAX™ supplement (Thermo Fisher Scientific, 11559726), 20ng/ml FGF2 (EUBIO, #450-33), 20ng/ml BMP4 (Bio-Techne Ltd. 5020-BP-010), N2.1 (Molecular Biology Service, VBC), Penicillin-Streptomycin (Sigma-Aldrich, P4333), 5μg/ml Heparin sodium salt from porcine intestinal mucosa (Sigma-Aldrich, H3393) and 2μg/ml Laminin (Sigma-Aldrich, L2020). Media changes were performed every 24h until D4 (Fig. 2 A, Fig. 3 A). A steady-state of quiescence was assumed after 72h of BMP4 treatment ^26,48^. Active NSCs were plated (15.000cells/cm²) in a separate day (D3, Fig 2 A) to avoid overgrow. Sampling started at D4 with collection every 4h spanning 36h (Immunostainings) or 48h (Sequencing). **Sequencing:** Cells were detached from the flask using Accutase (Sigma-Aldrich, A6964) and then collected in RNAse free DNA loBind tubes (Eppendorf, 22431021) using cold PBS. They were centrifuged at 5000 rpm in 4°C for 5 minutes. The supernatant was then removed carefully, and the pellet was kept at -70°C for further RNA extraction. **Coverslips:** Media was removed and carefully washed with 1X PBS. Cells were then fixed using 4% PFA (Sigma, cat.: P5148) washed once with 1X PBS and stored in PBS with 0.02% Sodium Azide (ITW Reagents, A1430) at 4°C for further processing.

### 4.7 Immunostaining and Image Analysis -*in vitro*

Blocking, primary antibody, and secondary antibody incubation conditions were identical to those used for brain sections, with the exception that the blocking and antibody solutions contained 0.1% Triton X-100. Coverslips were mounted using ProLong™ Glass Antifade Mountant (Thermo Fisher Scientific, cat. no. P36984). Images were acquired on the same microscope system as described above (**4.2**). Z-stacks were collected using a 40×/0.75 NA air objective (Olympus), spanning 10 µm of the specimen in 0.5 µm steps. A sum-of-slices Z-projection was applied, followed by automated nuclear segmentation using StarDist (fluorescence nucleus pre-trained model^49^). Mean gray values were extracted from the resulting nuclear masks. Rhythmicity was assessed by fitting a mixed-model cosinor with a fixed period of 24 h.

### 4.8 RNA Extraction

Total RNA was extracted using an in-house RNA isolation kit (VBCF, Vienna) based on guanidine thiocyanate lysis^50^, followed by magnetic bead-based purification (GE Healthcare, cat. no. 65152105050450) on a KingFisher automated sample purification system (Thermo Fisher Scientific, cat. no. 11859520).

### 4.9 Library Preparation and Sequencing

RNA quality and concentration were assessed using the Agilent DNF-471 SS Total RNA Kit on an Agilent Fragment Analyzer (Agilent Technologies). Only samples meeting minimum RNA integrity and concentration thresholds were used for downstream library preparation. Polyadenylated [poly(A)] RNA was enriched from total RNA using the NEBNext® Poly(A) mRNA Magnetic Isolation Module (New England Biolabs, cat. no. E7490), followed by strand-specific library construction with the NEBNext® Ultra II Directional RNA Library Prep Kit for Illumina® (New England Biolabs, cat. no. E7765), according to the manufacturer’s instructions. Library fragment size was assessed using the NGS High Sensitivity DNF-474-33 Kit on an Agilent Fragment Analyzer, and library concentrations were quantified using the KAPA Library Quantification Kit (Roche, cat. no. KK4824 / 07960140001). Sequencing was performed at the NGS Facility (Vienna BioCenter Core Facilities) on an Illumina NovaSeq X platform. Libraries were sequenced to a depth of 30 (± 5) million reads per sample.

### 4.10 RNAseq data processing and analysis

Raw reads were trimmed using Trimmomatic (v0.39) and subsequently aligned to the *Mus musculus* reference genome (GRCm39) using STAR (v2.7.11b). Gene-level read counts were quantified using featureCounts (Subread, v2.0.8). A DESeqDataSet was constructed from the gene-level count matrix and sample metadata using DESeq2, with a design formula including timepoint, experimental condition, and genotype (∼ timepoint + condition + genotype). To reduce the influence of lowly expressed genes, genes were retained if they had ≥10 raw counts in at least 39 samples, corresponding to the smallest group size in the experimental design. DESeq2 estimated size factors to normalize for differences in sequencing depth and library composition. Variance-stabilizing transformation (VST) was then applied to the raw count data using the vst function with blind = FALSE, incorporating the estimated dispersion trend while preserving the experimental design during transformation. The resulting VST-transformed expression matrix was used for downstream analyses, including dimensionality reduction and other exploratory analyses.

### 4.11 Detection of rhythmic transcripts in RNA-seq data

To detect rhythmicity, we employed a consensus approach followed by empirical validation through permutation testing. We applied ECHO^27^ and RAIN^28^ independently to each dataset (Active SGZ WT, Quiescent SGZ WT, Active SGZ P2KO, Quiescent SGZ P2KO, Active SVZ WT, and Quiescent SVZ WT) with a significance cutoff of p < 0.025. Transcripts were considered rhythmic only if detected by both methods. Additionally, transcripts classified by ECHO as “Overexpressed” or “Repressed” were discarded, retaining only those with Harmonic, Damped, or Forced oscillation profiles.

To control for spurious rhythmicity, a gene-specific empirical null was generated by permutation (1,000 iterations per condition). In each permutation, the temporal order of the timepoint blocks was shuffled independently for each gene, preserving the three-replicate structure within each timepoint while destroying the temporal (circadian) ordering. Each permuted matrix was passed through the identical RAIN→ECHO pipeline described above (same parameters, same p < 0.025 thresholds), and the number of permutations in which each gene was called rhythmic was recorded. For each gene, the proportion of permutations in which it was detected as rhythmic defined its empirical false-positive rate. Genes were retained as bona fide rhythmic only if this proportion was < 0.05, i.e. they were reproduced by chance in fewer than 5% of gene-specific null permutations, in addition to passing the RAIN and ECHO cutoffs in the observed data. This permutation-based approach was used in place of a parametric Benjamini-Hochberg correction because the RAIN null p-value distribution was markedly bimodal violating the uniform-null assumption on which parametric FDR control depends; the empirical filter makes no distributional assumption and yields a directly calibrated, gene-specific false-positive rate.

### 4.12 Soft Clustering Analysis and Enrichment Analysis

Fuzzy c-means clustering was performed using the *Mfuzz* R package to identify co-expressed gene clusters among rhythmic transcripts unique to each genotype. Analyses were conducted separately for active and quiescent SGZ-derived NSC datasets. VST-normalized counts were averaged across biological replicates per timepoint and standardized using the standardise function. The fuzzification parameter *m* was estimated for each dataset using the *mestimate* function. The optimal number of clusters was determined by evaluating within-cluster variation (*D*) across a range of 2–12 clusters using the Dmin function (3 repeats), and 8 clusters were selected for both datasets. Clustering was performed using the mfuzz function with the dataset-specific *m* parameter.

GO enrichment analysis was performed separately for each dataset using the *clusterProfiler* R package, with gene symbols converted to Entrez IDs via the bitr function using the org.Mm.eg.db annotation database. Enrichment was assessed for Biological Process (BP) ontology terms using the *enrichGO* function, with FDR adjustment and thresholds of p.adj < 0.05 and q < 0.2. Gene annotations were retrieved from Ensembl (v109) via *biomaRt*.

### 4.13 Differential Expression Analysis

We performed separate differential expression analyses on active and quiescent cell populations. Briefly, samples were subset from the merged dataset by cell state, and independent *DESeq2* models were fitted using a design formula incorporating timepoint as a covariate to control for temporal variance, and genotype as the primary variable of interest (∼timepoint + genotype). WT was set as the reference level for the genotype comparison. DEGs are defined as genes with an adjusted p-value < 0.05 and |log2 fold-change| ≥ 1.

### 4.14 Primary antibodies

4.14 Primary antibodies:
| Target molecule | Species | dilution | Figure | Company | Catalogue |
| --- | --- | --- | --- | --- | --- |
| **GFAP** | Mouse | 1:1000 | Fig. 1 (G-J)<br>Fig. 4<br>Fig. 5 (J-O) | SIGMA | G6171-200 |
| **GFAP** | Rat | 1:800 | Fig. 1 (C-F) | Invitrogen | 130300 |
| **SOX2** | Mouse | 1:50 | Fig. 1 (G-J)<br>Fig. 4<br>Fig. 5 (J-O) | Santa Cruz Biotechnology | sc-365964 |
| **SOX2** | Rat | 1:400 | Fig. 1 (C-F) | Invitrogen | 14-9811-82 / 14-9811-37 |
| **Ki67** | Rat | 1:200 | Fig. 4<br>Fig. 5 (J-O) | eBioscience/Invitrogen | 14-5698-82 |
| **BMAL1** | Rabbit | 1:1000 | Fig. 1 G–J, Fig. 2, Supp. Fig. 3– 4) | LeSauter et al., 2012 | n/a |
| **GFP** | Chicken | 1:5000 | Fig. 1 (C-F) | Abcam | ab13970 |
| **ASCL1** | Rabbit | 1:100 | Fig. 4 (B & D) | Abcam | ab211327 |
| **CCND1** | Rabbit | 1:100 | Fig. 4 (F & H) | Cell Signaling | 55506S |
| **MCM2** | Rabbit | 1:200 | Supp. Fig. 1-2 | Cell Signaling | 4007S |

### 4.15 Secondary antibodies

4. 15 Secondary antibodies:
| Target molecule | Conjugated fluorophore | Dilution | Company | Catalogue |
| --- | --- | --- | --- | --- |
| **Rabbit IgG** | Cy3 | 1:500 | Jackson Laboratories | 711-166-152 |
| **Rabbit IgG** | 647 | 1:500 | Jackson Laboratories | 711-606-152 |
| **Rat IgG** | 647 | 1:500 | Jackson Laboratories | 712-605-150 |
| **Chicken IgG** | 488 | 1:500 | Jackson Laboratories | 703-545-155 |
| **Mouse IgG** | 488 | 1:1000 | Invitrogen | A-21202 |
| **Mouse IgG** | 647 | 1:500 | Jackson Laboratories | 715-606-151 |

### 4.16 Data Availability

RNA-seq data are available on ENA under accession number PRJEB122945. Analysis code is available at github.com/pedrobrum1/CircadianAnalysis. Immunofluorescence quantifications are included in both the GitHub repository (*in vitro*) and in Supp. Table 1 (*in vivo*).

## References

1. Partch, C. L., Green, C. B. & Takahashi, J. S. Molecular architecture of the mammalian circadian clock. Trends in Cell Biology 24, 90–99 (2014).

2. Zhang, R., Lahens, N. F., Ballance, H. I., Hughes, M. E. & Hogenesch, J. B. A circadian gene expression atlas in mammals: Implications for biology and medicine. Proc. Natl. Acad. Sci. U.S.A. 111, 16219–16224 (2014).

3. Otobe, Y. et al. A mouse circadian proteome atlas. Molecular Cell 86, 393–406.e3 (2026).

4. Feillet, C., et al. Phase locking and multiple oscillating attractors for the coupled mammalian clock and cell cycle. Proc. Natl. Acad. Sci. U.S.A. 111, 9828–9833 (2014).

5. Farshadi, E., Van Der Horst, G. T. J. & Chaves, I. Molecular Links between the Circadian Clock and the Cell Cycle. Journal of Molecular Biology 432, 3515–3524 (2020).

6. Chakrabarti, S. et al. Hidden heterogeneity and circadian-controlled cell fate inferred from single cell lineages. Nat Commun 9, 5372 (2018).

7. Zhang, Z.-B., Sinha, J., Bahrami-Nejad, Z. & Teruel, M. N. The circadian clock mediates daily bursts of cell differentiation by periodically restricting cell-differentiation commitment. Proc. Natl. Acad. Sci. U.S.A. 119, e2204470119 (2022).

8. Brown, S. A. Circadian clock-mediated control of stem cell division and differentiation: beyond night and day. Development 141, 3105–3111 (2014).

9. Benitah, S. A. & Welz, P.-S. Circadian Regulation of Adult Stem Cell Homeostasis and Aging. Cell Stem Cell 26, 817–831 (2020).

10. Ming, G. & Song, H. Adult Neurogenesis in the Mammalian Brain: Significant Answers and Significant Questions. Neuron 70, 687–702 (2011).

11. Urbán, N., Blomfield, I. M. & Guillemot, F. Quiescence of Adult Mammalian Neural Stem Cells: A Highly Regulated Rest. Neuron 104, 834–848 (2019).

12. Bottes, S. et al. Long-term self-renewing stem cells in the adult mouse hippocampus identified by intravital imaging. Nat Neurosci 24, 225–233 (2021).

13. Encinas, J. M. et al. Division-Coupled Astrocytic Differentiation and Age-Related Depletion of Neural Stem Cells in the Adult Hippocampus. Cell Stem Cell 8, 566–579 (2011).

14. Borgs, L. et al. Period 2 regulates neural stem/progenitor cell proliferation in the adult hippocampus. BMC Neurosci 10, 30 (2009).

15. Bouchard-Cannon, P., Mendoza-Viveros, L., Yuen, A., Kærn, M. & Cheng, H.-Y. M. The Circadian Molecular Clock Regulates Adult Hippocampal Neurogenesis by Controlling the Timing of Cell-Cycle Entry and Exit. Cell Reports 5, 961–973 (2013).

16. Gengatharan, A. et al. Adult neural stem cell activation in mice is regulated by the day/night cycle and intracellular calcium dynamics. Cell 184, 709–722.e13 (2021).

17. Liu, Q., et al. Coordination between circadian neural circuit and intracellular molecular clock ensures rhythmic activation of adult neural stem cells. Proc. Natl. Acad. Sci. U.S.A. 121, e2318030121 (2024).

18. Ebihara, S., Marks, T., Hudson, D. J. & Menaker, M. Genetic Control of Melatonin Synthesis in the Pineal Gland of the Mouse. Science 231, 491–493 (1986).

19. Kasahara, T., Abe, K., Mekada, K., Yoshiki, A. & Kato, T. Genetic variation of melatonin productivity in laboratory mice under domestication. Proc. Natl. Acad. Sci. U.S.A. 107, 6412– 6417 (2010).

20. Huang, S. et al. Demyelination Regulates the Circadian Transcription Factor BMAL1 to Signal Adult Neural Stem Cells to Initiate Oligodendrogenesis. Cell Reports 33, 108394 (2020).

21. Smyllie, N. J. et al. Visualizing and Quantifying Intracellular Behavior and Abundance of the Core Circadian Clock Protein PERIOD2. Current Biology 26, 1880–1886 (2016).

22. LeSauter, J. et al. Antibodies for Assessing Circadian Clock Proteins in the Rodent Suprachiasmatic Nucleus. PLoS ONE 7, e35938 (2012).

23. Mirzadeh, Z., Merkle, F. T., Soriano-Navarro, M., Garcia-Verdugo, J. M. & Alvarez-Buylla, A. Neural Stem Cells Confer Unique Pinwheel Architecture to the Ventricular Surface in Neurogenic Regions of the Adult Brain. Cell Stem Cell 3, 265–278 (2008).

24. Codega, P., et al. Prospective Identification and Purification of Quiescent Adult Neural Stem Cells from Their In Vivo Niche. Neuron 82, 545–559 (2014).

25. Gabarró-Solanas, R. et al. Adult neural stem cells and neurogenesis are resilient to intermittent fasting. EMBO Rep 24, EMBR202357268 (2023).

26. Mira, H. et al. Signaling through BMPR-IA Regulates Quiescence and Long-Term Activity of Neural Stem Cells in the Adult Hippocampus. Cell Stem Cell 7, 78–89 (2010).

27. De Los Santos, H. et al. ECHO: an application for detection and analysis of oscillators identifies metabolic regulation on genome-wide circadian output. Bioinformatics 36, 773–781 (2020).

28. Thaben, P. F. & Westermark, P. O. Detecting Rhythms in Time Series with RAIN. J Biol Rhythms 29, 391–400 (2014).

29. Bae, K. et al. Differential Functions of mPer1, mPer2, and mPer3 in the SCN Circadian Clock. Neuron 30, 525–536 (2001).

30. Urbán, N. et al. Return to quiescence of mouse neural stem cells by degradation of a proactivation protein. Science 353, 292–295 (2016).

31. Imayoshi, I. et al. Oscillatory Control of Factors Determining Multipotency and Fate in Mouse Neural Progenitors. Science 342, 1203–1208 (2013).

32. Miller, I. et al. Ki67 is a Graded Rather than a Binary Marker of Proliferation versus Quiescence. Cell Reports 24, 1105–1112.e5 (2018).

33. Janich, P. et al. The circadian molecular clock creates epidermal stem cell heterogeneity. Nature 480, 209–214 (2011).

34. Mortimer, T. et al. The epidermal circadian clock integrates and subverts brain signals to guarantee skin homeostasis. Cell Stem Cell 31, 834–849.e4 (2024).

35. Matsu-ura, T. et al. Intercellular Coupling of the Cell Cycle and Circadian Clock in Adult Stem Cell Culture. Molecular Cell 64, 900–912 (2016).

36. Schouten, M. et al. Circadian glucocorticoid oscillations preserve a population of adult hippocampal neural stem cells in the aging brain. Mol Psychiatry 25, 1382–1405 (2020).

37. Silva-Vargas, V., Maldonado-Soto, A. R., Mizrak, D., Codega, P. & Doetsch, F. Age-Dependent Niche Signals from the Choroid Plexus Regulate Adult Neural Stem Cells. Cell Stem Cell 19, 643–652 (2016).

38. Fame, R. M. et al. Defining diurnal fluctuations in mouse choroid plexus and CSF at high molecular, spatial, and temporal resolution. Nat Commun 14, 3720 (2023).

39. Leeman, D. S. et al. Lysosome activation clears aggregates and enhances quiescent neural stem cell activation during aging. Science 359, 1277–1283 (2018).

40. Knobloch, M. et al. Metabolic control of adult neural stem cell activity by Fasn-dependent lipogenesis. Nature 493, 226–230 (2013).

41. Kobayashi, T. et al. Enhanced lysosomal degradation maintains the quiescent state of neural stem cells. Nat Commun 10, 5446 (2019).

42. O’Neill, J. S. & Reddy, A. B. Circadian clocks in human red blood cells. Nature 469, 498–503 (2011).

43. Ch, R. et al. Rhythmic glucose metabolism regulates the redox circadian clockwork in human red blood cells. Nat Commun 12, 377 (2021).

44. Haydon, M. J., Hearn, T. J., Bell, L. J., Hannah, M. A. & Webb, A. A. R. Metabolic regulation of circadian clocks. Seminars in Cell & Developmental Biology 24, 414–421 (2013).

45. Taufique, S. K. T. et al. A novel circadian behavior rhythm in arrhythmic Period triple knockout mice revealed by constant light exposure. iScience 28, 113292 (2025).

46. Schindelin, J. et al. Fiji: an open-source platform for biological-image analysis. Nat Methods 9, 676–682 (2012).

47. Hagihara, H., Toyama, K., Yamasaki, N. & Miyakawa, T. Dissection of Hippocampal Dentate Gyrus from Adult Mouse. JoVE 1543 (2009) doi:10.3791/1543.

48. Blomfield, I. M. et al. Id4 promotes the elimination of the pro-activation factor Ascl1 to maintain quiescence of adult hippocampal stem cells. eLife 8, e48561 (2019).

49. Schmidt, U., Weigert, M., Broaddus, C. & Myers, G. Cell Detection with Star-Convex Polygons. in Medical Image Computing and Computer Assisted Intervention – MICCAI 2018 (eds Frangi, A. F., Schnabel, J. A., Davatzikos, C., Alberola-López, C. & Fichtinger, G.) vol. 11071 265–273 (Springer International Publishing, Cham, 2018).

50. Boom, R. et al. Rapid and simple method for purification of nucleic acids. J Clin Microbiol 28, 495–503 (1990).

